# Targeted glycophagy ATG8 therapy reverses diabetic heart disease in mice and in human engineered cardiac tissues

**DOI:** 10.1101/2024.11.28.625926

**Authors:** K M. Mellor, U. Varma, P. Koutsifeli, C.L. Curl, J.V. Janssens, L.J. Daniels, G.B. Bernasochi, A.J.A. Raaijmakers, M. Annandale, X. Li, S.L. James, D.J. Taylor, K. Raedschelders, K.L. Weeks, R.J. Mills, R.G. Parton, X. Hu, J.R. Bell, Terence J. O’Brien, Rajesh Katare, E.R. Porrello, J.E. Hudson, R-P. Xiao, J.E. Van Eyk, R.A. Gottlieb, L.M.D. Delbridge

## Abstract

Diabetic heart disease is highly prevalent and is associated with the early development of impaired diastolic relaxation. The mechanisms of diabetic heart disease are poorly understood and it is a condition for which there are no targeted therapies. Recently, disrupted glycogen-autophagy (glycophagy) and glycogen accumulation have been identified in the diabetic heart. Glycophagy involves glycogen receptor binding and linking with an ATG8 protein to locate and degrade glycogen within an intracellular phago-lysosome. Here we show that glycogen receptor protein STBD1 (starch-binding-domain-protein-1) is mobilized early in the cardiac glycogen response to metabolic challenge *in vivo*, and that deficiency of a specific ATG8 family protein, Gabarapl1 (γ-aminobutyric-acid-receptor-associated-protein-like-1) is associated with diastolic dysfunction in diabetes. Gabarapl1 gene delivery treatment remediated cardiomyocyte and cardiac diastolic dysfunction in type 2 diabetic mice and diastolic performance of ‘diabetic’ human iPSC-derived cardiac organoids. We identify glycophagy dysregulation as a mechanism and potential treatment target for diabetic heart disease.

## MAIN TEXT

Diabetic heart disease is recognized as a distinct cardiomyopathy, characterized by occurrence of diastolic dysfunction.^1^ Whilst derangements of cardiomyocyte signaling and functional performance have been described in various forms of diabetic cardiopathology (including types 1 and 2 diabetes),^5,6^ the underlying mechanisms which drive the development of diastolic dysfunction and progression to heart failure in diabetes are unclear.

Autophagy is a highly conserved essential catabolic process which involves the breakdown and ‘recycling’ of defective or surplus cellular materials. An autophagy role in metabolic disease-related heart pathology was first reported more than a decade ago^7,8^ describing altered levels of an autophagy intermediate protein in an experimental model of type 2 diabetes. In overview, autophagic processes include receptor tagging of ‘cargo’ destined for intracellular breakdown. Receptors link with membrane-bound ATG8 family proteins and recruit the molecular machinery to enclose the cargo into an autophagosome. Lysosome fusion with the phagosome allows breakdown and phago-lysosome export of cargo content.^3^

It is now clear that various autophagy receptors and associated ATG8 partners can target specific cellular components. Work in non-muscle cell culture lines has linked a specific glycogen receptor STBD1 (Starch binding domain protein 1) with an ATG8 partner GABARAPL1 (γ GABA type A receptor associated protein like 1).^4,9^ Evidence demonstrates that glycogen-selective autophagy (‘glycophagy’) is operational in the myocardium. Our studies in primary cardiomyocyte cultures have shown STBD1 localization to be distinct and separate to the more extensively characterized ATG8 protein LC3B,^10^ which mediates autophagy of protein aggregates via the p62 receptor partner. It has also been shown *in vitro* that cardiomyocyte cellular glycogen levels respond to culture media simulating diabetic hyperglycemic conditions.^10^

As a dense hexose sugar polymer, glycogen constitutes a concentrated glycolysis fuel store. Cardiac glycogen is considered an important energy reserve in physiological settings.^11^ Glycolysis is a key energy source in cardiac electro-mechanical transduction, particularly involved in supporting SERCA2-mediated Ca^2+^ reuptake during diastolic relaxation.^12,13^ We have recently shown that marked increases in glycogen levels are a consistent feature in a range of diabetic pre-clinical models and in clinically-derived cardiac tissues.^2^ This as yet unexplained phenomenon is cardiac specific – skeletal muscle glycogen levels are not perturbed in basal diabetic pathologic states.^2^ Furthermore, cardiac glycogen elevation in rodent diabetic myocardium is associated with electron microscopic visualization of ectopically clustered glycogen particulates amassed adjacent to mitochondria and dispersed throughout the myofilament architecture. In these settings, we have demonstrated that formed and forming glycogen-enriched phagosomes are prominent structures, suggesting cardiomyocyte glycophagy disruption in diabetes.^2^ Our recent *in vivo* and *in vitro* studies have shown that myocardial glycophagy flux (i.e. autophagic glycogen throughput) is impaired in diabetes.^2,14^

Thus, in parallel with the well characterized cytosolic enzymatic regulation of glycogen synthesis and breakdown (by glycogen synthase and phosphorylase), glycophagy constitutes a route for ‘bulk’ homeostatic glycogen handling,^4^ susceptible to disruption in diabetes. We hypothesize that in diabetes, disturbance of glycogen processing through glycophagy contributes to cardiac dysfunction, in particular impairing relaxation and diastolic performance. To directly test this proposition, in the present study we sought to determine whether *in vivo* gene manipulation of the key glycophagy ATG8 intermediate (GABARAPL1), has therapeutic potential in diabetic heart disease.

### Glycophagy mediator proteins are mobilized early in the cardiac glycogen response to metabolic challenge *in vivo*

Firstly, we used proteomic methods (LC-MS/MS) to explore the cardiomyocyte glycogen receptor - ATG8 homolog relationship, to confirm previous *in vitro* immuno-characterization, and ensure the most efficacious gene targeting approach for glycophagy intervention. STBD1 is known to bind to glycogen at a carbohydrate binding module site (CBM20), and also contains several ATG8-interacting motifs (AIMs). Our *in silico* analyses revealed that 12 glycogen-related proteins contain AIMs (including STBD1) and therefore have the potential to be glycogen receptors to partner with ATG8 proteins.^4^ There are 6 ATG8 homologues: the microtubule-associated protein 1A/1B-light chain 3 series (LC3A, LC3B, LC3C); and the γ-aminobutyric acid receptor-associated protein /like1 series (GABARAP, GABARAPL1 and GABARAPL2). Thus potentially numerous receptor-ATG8 combinations are possible.

To contrast the cardiac- and skeletal muscle-specific glycogen proteomes, and to determine whether (and which) CBM20 domain proteins might be present at the earliest stage of systemic metabolic disturbance, an experimental acute metabolic stimulus was used. A metabolic challenge was induced in rats with a single dose of streptozotocin (STZ) to lesion pancreatic β cells to suppress insulin secretion. Although multi-dose STZ is commonly used to induce a model of type 1 diabetes, for our study the acute STZ intervention afforded an experimental setting of rapid onset metabolic challenge (MC) to interrogate the early glycogen response. After 48 hours, blood glucose was elevated to more than 20 mM in all treated animals (Ctrl: 6.5 ± 0.2 mM vs. MC: 24.6 ± 0.8 mM). We confirmed significant glycogen elevation in cardiac, but not skeletal muscle tissues (Fig. 1a). Thus cardiac glycogen accumulation was identified as a primary feature of cardiometabolic challenge, and a response evident early after systemic metabolic disturbance. Proteomic analyses were performed on glycogen extracts using liquid chromatography and mass spectrometry (LC-MS/MS). Overall, the cardiac glycogen proteome was more extensive than the skeletal glycogen proteome, comprising a larger number of total identified proteins (513 vs 366 proteins), indicative of increased complexity of the cardiac glycogen regulatory environment (Fig. 1b). For each tissue, amongst the common proteins detected in both control and challenge conditions, a relatively small number showed differential abundance (cardiac n=19, skeletal n=7). Notably, the response to metabolic challenge involved recruitment of unique components to the glycogen proteome in both tissues.

**Fig. 1.**
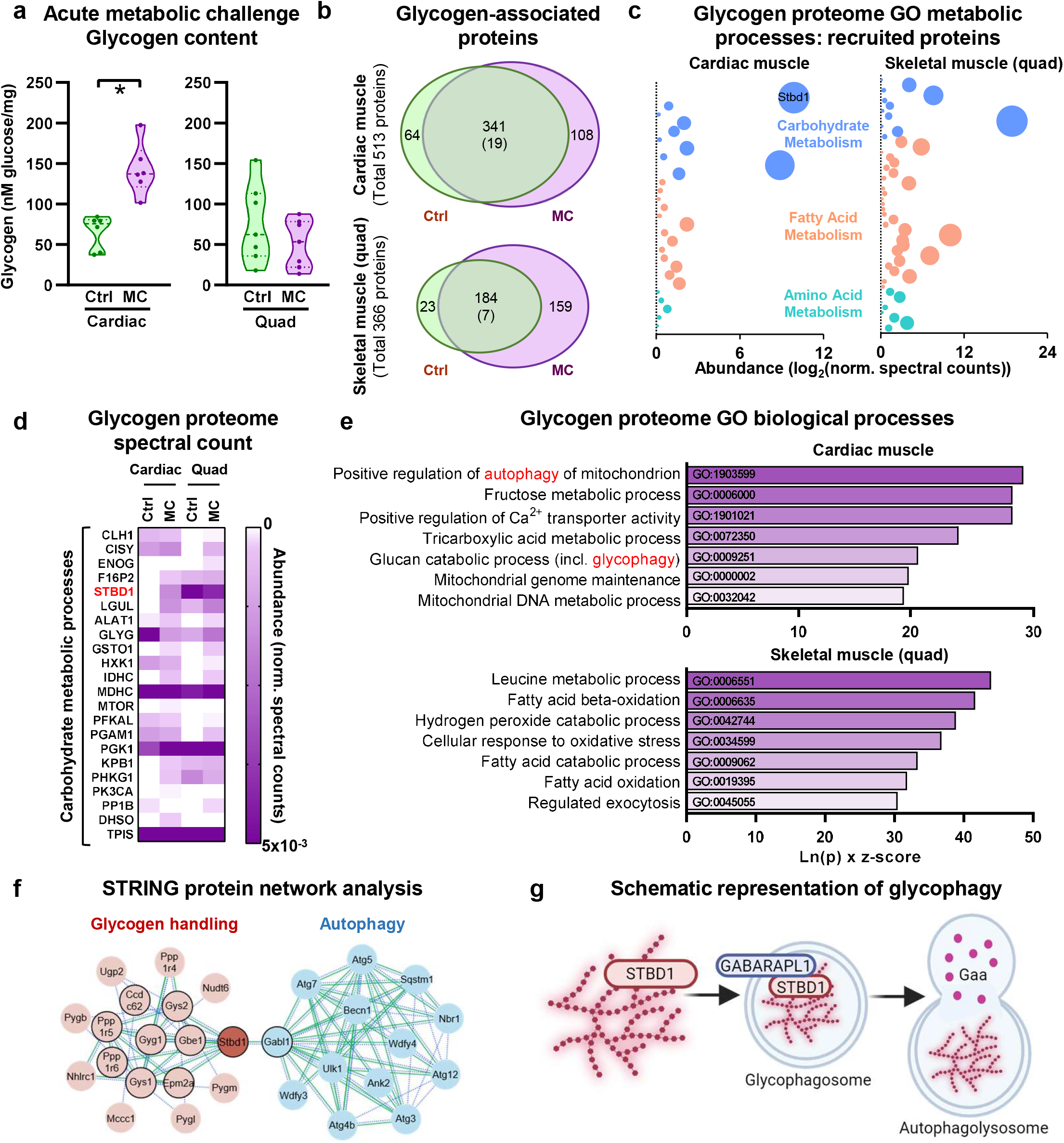
Glycophagy mediator proteins are mobilized early in cardiac glycogen response to metabolic challenge *in vivo*. **a**, Heart glycogen is increased in rats subjected to an acute metabolic challenge (MC, 48 hr post-streptozotocin injection) and skeletal muscle glycogen is unchanged (analyzed by unpaired 2-sided T-Test. Heart: n=6 animals/group, p=0.0006. Skeletal muscle: n=7 animals/group, p=0.28). **b**, Glycogen proteome in cardiac and skeletal (quadriceps) muscle response to an acute metabolic challenge *in vivo*. Venn diagram values show the number of unique or differentially abundant proteins in control (Ctrl) vs metabolic challenge (MC) samples using LC-MS/MS. Total number of significantly up- or down-regulated proteins shown in brackets. n=4 animals /group. **c**, The glycogen proteome response to an acute metabolic challenge is characterized by changes in proteins related to carbohydrate and amino acid metabolism in cardiac muscle, contrasting with skeletal muscle where changes in fatty acid metabolism related proteins are most evident. Data are derived from spectral counts of uniquely present proteins in acute metabolic challenge rat tissue from Carbohydrate (GO:0005975), Fatty Acid (GO:0006629) or Amino Acid (GO:0006520) Metabolic Process GO categories, detected by LC-MS/MS (n=4 animals/group). Bubble size is indicative of protein abundance. **d**, The glycophagy tagging protein, Stbd1 (starch-binding domain-containing protein 1) is detected as a component of the cardiac glycogen proteome only in acute metabolic challenge cardiac muscle (but not in control), whereas Stbd1 is present in both metabolic challenge and control skeletal muscle glycogen proteome. The heatmap is derived from LC-MS/MS normalized spectral counts of differentially abundant or uniquely detected proteins in the Carbohydrate Metabolic Processes GO category (GO:0005975), with normalized spectral abundance factor adjustment (n=4 animals /group). **e**, Glycophagy is identified as a key biological process in the cardiac glycogen proteome response to an acute metabolic challenge *in vivo*. The gene ontology (GO) categories with the highest combined score (Ln(p) x z-score) are presented, comprising glycogen proteome proteins uniquely detected or differentially abundant between metabolic challenge and control for heart and skeletal muscle (n=4 animals/group). **f**, Functional analysis network (STRING) of known and predicted protein interactions with Stbd1. Gabarapl1 is identified as a primary interactor with Stbd1 and a link between glycogen handling and autophagy. Primary interactions are depicted by black outline. Associations are derived from experimental evidence (continuous green line) and text mining evidence (dashed blue line). **g**, Schematic illustrating three stages of glycophagy. Stbd1 tags glycogen and binds to the Atg8 protein, Gabarapl1 (GABA receptor-associated protein-like 1), at the forming glycophagosome membrane. The mature glycophagosome fuses with a lysosome where Gaa (acid α-glucosidase) degrades glycogen to free glucose for metabolic recycling. Made with Biorender Data are presented as mean ± s.e.m. *p<0.05. See also Extended Data Figure 1 and Extended Data Table 1.

To explore the metabolic characteristics of these tissue-specific responses to metabolic challenge we searched the group of recruited proteins using Gene Ontology terms for Carbohydrate, Fatty Acid and Amino Acid Metabolic Processes (GO:0005975, GO:0006629, GO:0006520). The proteomic profiles generated for each tissue type were markedly different (Fig. 1c), with cardiac muscle proteins showing carbohydrate protein dominance, while fatty acid metabolism-related proteins were more prominent in skeletal muscle. Given that ‘lipotoxicity’ is a frequent descriptor used in relation to metabolic disturbance in cardiac muscle pathology,^15^ this finding was unexpected. These observations indicate that myocardial carbohydrate metabolic derangement has an ‘early responder’ role in cardiomyocyte glycemic dysregulation.

As an identified glycogen receptor, detection of STBD1 was expected.^4,9,16^ Remarkably it was the most abundant protein of the GO ‘Carbohydrate Metabolism’ term (annotated Fig. 1c), and was recruited to the cardiac proteome in response to metabolic challenge and not the skeletal proteome. Regarding the other 11 known glycogen-related proteins with AIM sequences, glycogen synthase 1 (GYS1), glycogen phosphorylase (PYGM, PYGL, PYGB), and laforin (EPM2A) were detected in cardiac extract but not differentially abundant between metabolic challenge and control. Glycogen synthase 2 (GYS2), glycogen branching enzyme (AGL), glycogenin 2 (GYG2), glycogen debranching enzyme (GDE), and malin (NHLRC1) were not detected in either extract. Glycogenin 1 (GLYG1) had 25% lower abundance in cardiac glycogen extracted from the metabolic challenge vs control rats. For the pooled group of glycogen-associated proteins uniquely recruited or differentially abundant in the ‘carbohydrate metabolic processes’ GO category, a map was constructed to compare cardiac and skeletal muscle proteomes (Fig. 1d, Extended Data Table 1). Amongst the 22 proteins detected, STBD1 abundance characteristics were distinctive – under control conditions STBD1 was robustly present in the skeletal glycogen proteome, but not detectable in the cardiac glycogen extracts. In response to the metabolic challenge, while STBD1 was recruited to the cardiac glycogen proteome, the abundance was simultaneously diminished in the skeletal glycogen proteome. The more vigorous cardiac glycogen response to metabolic challenge (vs skeletal muscle) is consistent with clinical findings that cardiac phenotype is most markedly disrupted in a range of somatic genetic conditions described as ‘glycogen storage diseases’.^17^

Next, a GO analysis was undertaken (Fig. 1e) to obtain information about higher order functional impacts of metabolic challenge for each tissue using the full sets of differentially abundant and uniquely detected proteins in both treatments (i.e. cardiac n = 64 + 19 + 108 = 191 proteins; skeletal n = 23 + 7 + 159 = 189 proteins (i.e. uniquely control + differentially abundant + uniquely MC)). The higher relative change in glycogen-associated proteins in skeletal muscle (52% vs 37%) may reflect a more complex adaptation to maintain glycogen stores in metabolic stress settings. The cardiac response was characterized by changes in autophagy and glucan catabolic processes in concert with mitochondrial involvement. In contrast, the highest ranked processes in the skeletal muscle response were fatty acid catabolism and oxidative stress. Thus, very different glycogen proteomic metabolic responses were observed in different tissues of the same animals exposed to *in vivo* systemic metabolic challenge induced by acute insulin depletion.

Collectively, these proteomic data obtained by targeting the earliest phase of systemic metabolic disturbance reveal that cardiac and skeletal glycogen ‘first responder’ processes are disparate. It may be inferred that glycogen is primarily used episodically relating to activity demand in skeletal muscle, while in cardiac muscle glycogen constitutes an important line of fuel defense/reserve as a response to metabolic stress. The cardiac glycogen proteome appears to be more nuanced and protein recruitment more aligned with carbohydrate metabolic adaptations, compared to skeletal muscle where glycogen levels are maintained and subtle shifts in fatty acid metabolism are observed in a setting of metabolic challenge. The cardiac metabolic response pattern in this acute setting, associated with elevation of myocardial glycogen, is consistent with the observed chronic state of glycogen accumulation in diabetic heart disease and suggests potential for disease causation. Augmented STBD1 recruitment to the cardiac glycogen proteome emerges as a crucial point of difference in determining the unique cardiac-patho-phenotype in the initial response to deranged systemic metabolism.

Based on the glycogen proteome data, we next performed a functional network analysis to probe the possible links between glycogen receptor and autophagosome ATG8 effector proteins in metabolic challenge. The STRING database (Search Tool for Retrieval of Interacting Genes/Proteins) captures experimental evidence and performs text-mining to identify evidence of direct and indirect protein-protein associations. Using STBD1 as the input protein, an association network was generated and depicted (Fig. 1f) to illustrate primary (direct, black outline) and secondary protein interactions and designated links of specific evidence type.

Using STRING analysis, GABARAPL1 emerged as the only autophagy protein with direct link to STBD1. This *in silico* finding is supported by several literature reports confirming STBD1-GABARAPL1 binding.^16,18,19^ Recent experimental work resolved the crystal structure of STBD1 interactions with ATG8 family proteins and showed that STBD1 binding affinity was strongest with GABARAPL1.^19^ It was also determined that the two oligosaccharide binding sites of STBD1 can (uniquely) accommodate glucose polymer structures involving both the α-1,4- and α-1,6-glycosidic bond types, corresponding to linear and branched glycogen configurations.^19^

Cumulatively these functional proteomic investigations confirm this receptor-Atg8 relationship to constitute a prominent cardiac glycogen autophagy axis. Based on these findings, and a growing general understanding of phagosome mechanisms,^3,4,20^ cardiac glycophagy can be conceptualized to comprise several stages: the initial STBD1 receptor attachment to glycogen (CBM20 domain); the glycogen-attached receptor linking to GABARAPL1 at the forming autophagosome membrane (via an ATG8-Interacting Motif, AIM); membrane enclosure of glycogen cargo, and subsequent fusion of the auto-phagosome with a lysosome (containing acid α-glucosidase, GAA) to degrade glycogen generating hexose sugar for cell recycling (Fig. 1g).

### Cardiac GabarapL1 (Atg8) deficiency is associated with diastolic dysfunction in diabetes

Next, we investigated the link between glycogen, glycophagy and diastolic dysfunction in established diabetes. Given the known cardiac contractile deficits associated with diabetes,^21^ we tested whether a functional association with cardiac glycogen elevation could be identified in a chronic type 2 diabetic (T2D) setting. A high fat (with high sugar) dietary treatment was used to provide a clinically-relevant T2D model of glucose intolerance, mild hyperglycemia, and obesity (Extended Data Table 2), opting for a more moderate hyperglycemia than can be evident in other rodent models of T2D (e.g. STZ, ZDF, db/db).^22^ This investigation acquired matching echo data derived *in vivo* with glycogen analysis of tissues recovered post-mortem. Cardiac glycogen content was elevated in T2D mice (Fig. 2a) in the absence of changes in glycogen synthase and glycogen phosphorylase (Extended Data Fig. 1a-b). Diastolic dysfunction *in vivo*, as typically indexed by elevated ratio of the mitral blood flow/tissue displacement Doppler signals (E/e’), was apparent in T2D mice (Fig. 2a, Extended Data Fig. 1c). The elevated E/e’ levels recorded were proportionally commensurate with shifts reported in patients with type 2 diabetes.^23^ A significant correlation between glycogen and diastolic dysfunction was demonstrated (Fig. 2a). No evidence of altered systolic function (including ejection fraction, fractional shortening) was detected (Extended Data Fig. 1c-d). A significant correlation between glycogen and diastolic dysfunction was also evident in isolated cardiomyocytes from chronic STZ rats. Elevated cardiomyocyte glycogen content was correlated with prolonged cardiomyocyte Ca^2+^ transient decay (Tau, Extended Fig. 1e-g). Given our previous report that cardiac glycophagy flux is impaired in diabetes,^2^ we evaluated the expression level of the key glycophagy-mediator genes, *Stbd1, Gabarapl1* and *Gaa* to identify the optimal target for gene therapy intervention.

**Fig. 2.**
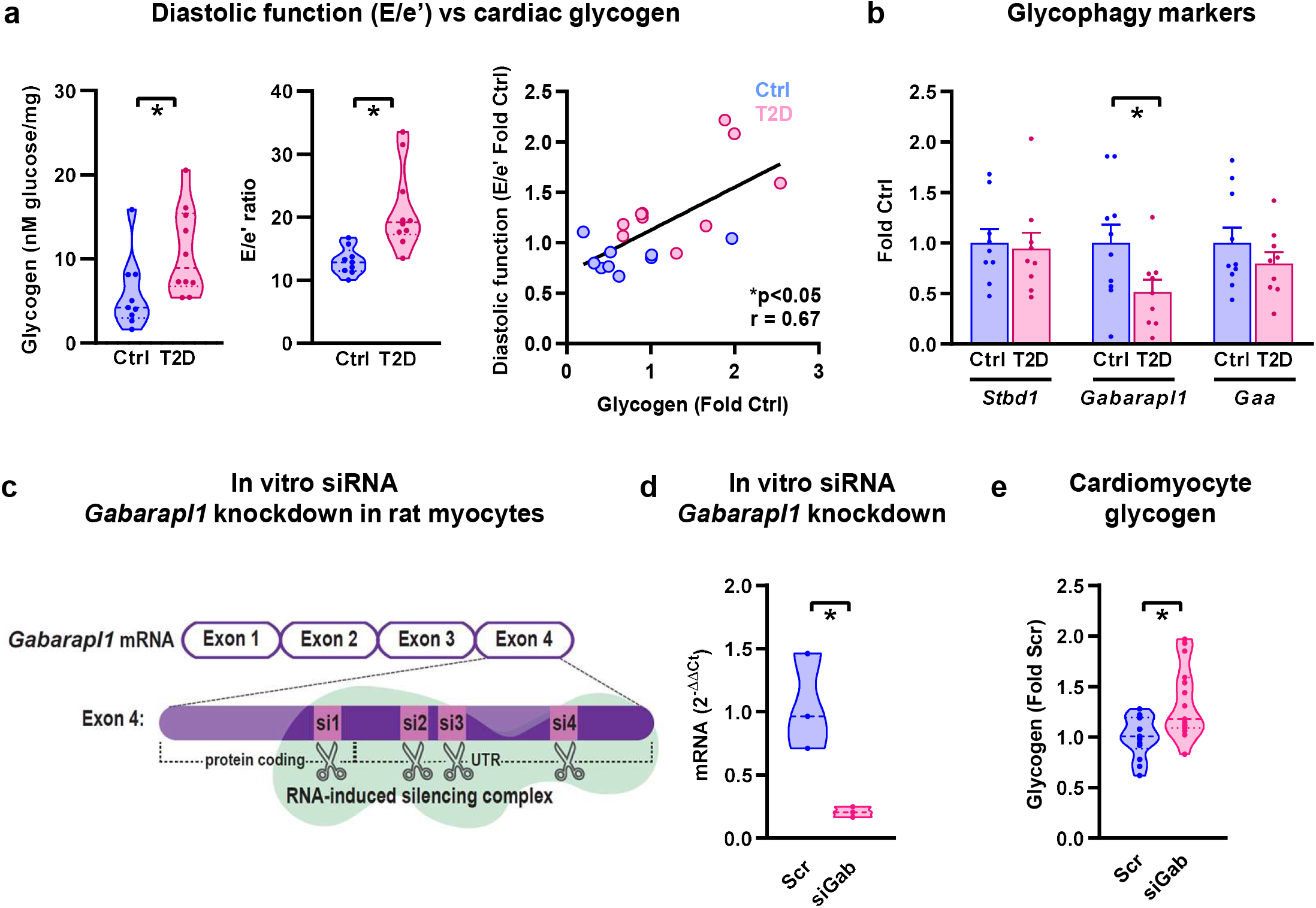
Cardiac Gabarapl1 (Atg8) deficiency is associated with diastolic dysfunction in diabetes *in vivo* and induces glycogen accumulation *in vitro*. **a**, Diastolic dysfunction (increased echocardiographic index, E/e’ ratio, p=0.0024) and myocardial glycogen accumulation (p=0.0397) are evident in type 2 diabetic mice (T2D, high fat diet, unpaired 2-sided T-Test). Myocardial glycogen content and diastolic dysfunction (E/e’ ratio) are correlated for control (blue) and T2D (pink) mice (r, Pearson correlation coefficient, Ctrl: n=9 and T2D: n=10 animals/group). **b**, Cardiac mRNA expression of glycophagy markers in T2D mice. Gabarapl1 is decreased (p=0.0045) and Stbd1 and Gaa mRNA are unchanged (Ctrl Gabarapl1, Gaa: n=10 animals/group, Ctrl Stbd1, T2D Stbd1, Gabarapl1 & Gaa: n=9 animals/group, unpaired 2-sided T-Test). **c**, Schematic depicting small interfering RNA (siRNA) gene silencing experimental design using a pool of 4 siRNA sequences (si1-4) targeting Gabarapl1 for gene knockdown in neonatal rat ventricular myocytes (NRVM). **d**, Confirmation of siRNA-induced Gabarapl1 mRNA knockdown (siGab) in NRVMs (n=3 independent wells, p=0.019, unpaired 2-sided T-test). **e**, Cardiomyocyte glycogen content increased in NRVMs in response to Gabarapl1 knockdown (Scr: n=14, siGab: n=15 wells from 3 biologically independent cell culture experiments, p=0.0036, unpaired 2-sided T-test). Data are presented as mean ± SEM. *p<0.05. See also Extended Data Figure 2 and Extended Data Table 2,3,&4.

Gene expression levels of *Stbd1* and *Gaa* were preserved in T2D mice. In contrast, a dramatic 60% reduction in *Gabarapl1* mRNA was observed (Fig. 2b, Extended Data Table 3), suggesting that ATG8 deficiency may be a glycophagy flux limiting factor. In clinical atrial biopsies from patients with diabetes a similar finding was observed, although the data were more variable and the extent of difference was more modest. Lower *Gabarapl1* mRNA expression and a trend for lower GABARAPL1 protein in the membrane:cytosol fractions and accumulation of STBD1 protein were observed in patients with diabetes (Extended Data Fig. 1h-j, Extended Data Table 4). To further establish the link between Gabarapl1 and diastolic dysfunction in a controlled in vitro setting, we used a gene knockdown approach. *In vitro* knockdown of *Gabarapl1* in cultured neonatal rat ventricular myocytes using siRNA treatment produced a marked increase in cardiomyocyte glycogen content (Fig. 2c-e, Extended Data Table 3). *In vivo* knockdown of *Gabarapl1* using a Crispr/Cas9 gene targeting approach was pursued to generate heterozygote Gabarapl1-KO mice (Fig. 3a, Extended Data Fig. 2a-d, Extended Data Table 3). Gabarapl1-KO mice (age 10 weeks) exhibited normal systemic characteristics with no change in body weight, glucose tolerance and blood glucose levels (Fig. 3b-d). Diastolic dysfunction was evident in Gabarapl1-KO mice, evidenced by increased E/e’ with preservation of systolic function (ejection fraction, fractional shortening; Fig. 3e-g). The original and seminal report of cardiac autophagy^24^ identified the neonatal period of nutrition transition to be particularly reliant on autophagy activity. Thus, to assess the impact of Gabarapl1 deficiency on myocardial glycogen levels, tissues of adult and neonate (age 2 days) mice were examined. Evidence of glycogen accumulation was observed at both ages (Fig. 3h,i).

**Fig. 3.**
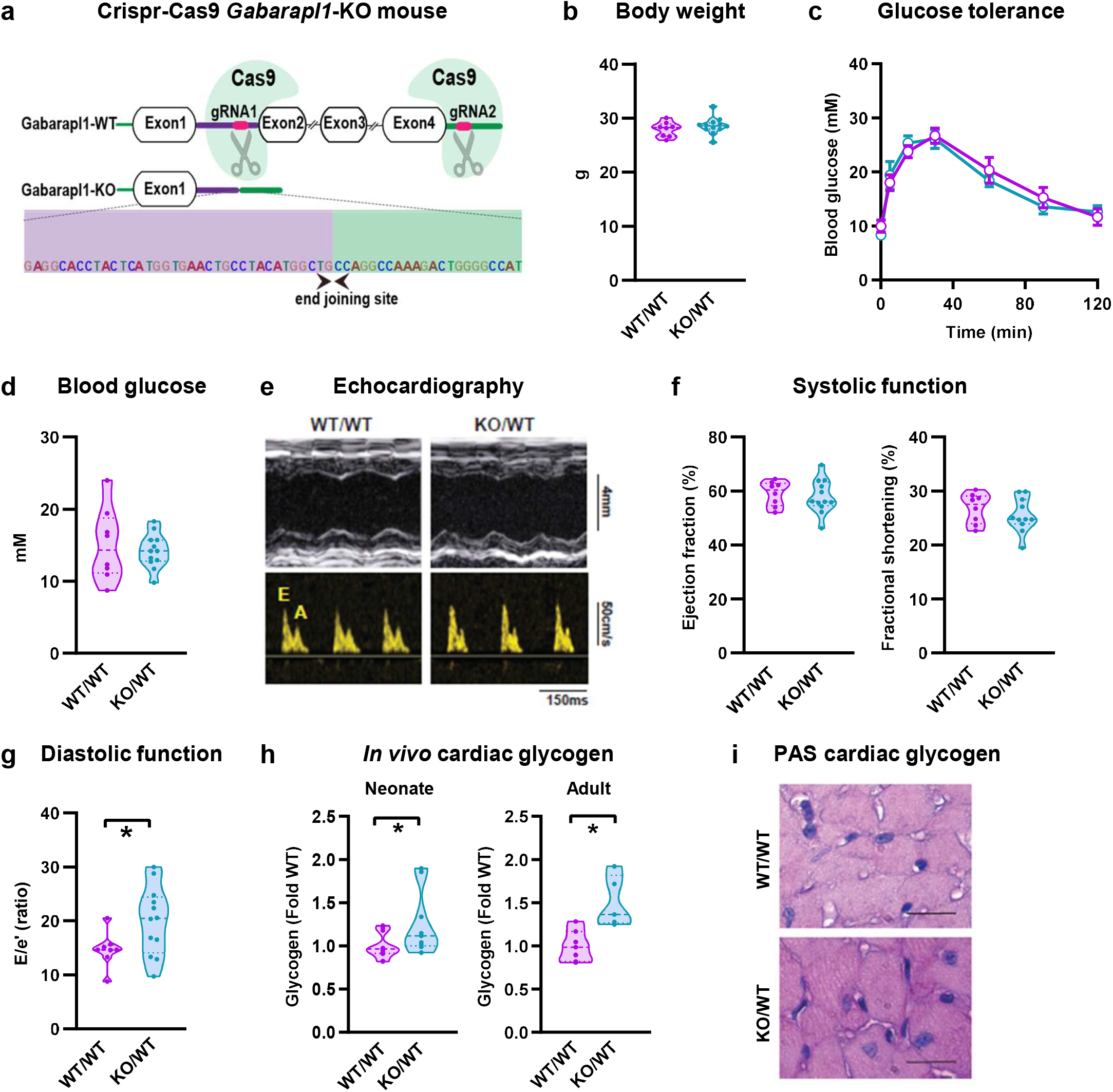
Gabarapl1 knockdown induces diastolic dysfunction and cardiac glycogen accumulation in mice. **a**, Schematic of Crispr/Cas9 genome editing design targeting *Gabarapl1* for gene deletion in mice. Global *Gabarapl1* knockdown did not affect **b**, body weight (WT: n=8, KO/WT: n=12 animals/group, unpaired 2-sided T-test), **c**, glucose tolerance (WT: n=3, KO/WT: n=7 animals/group, repeated measures ANOVA with Bonferroni post-hoc), or **d**, blood glucose levels (WT: n=8, KO/WT: n=12 animals/group, unpaired 2-sided T-test). Despite no effect on systemic parameters, a cardiac phenotype was evident. **e**, M-mode echocardiography exemplar traces from left ventricular short axis view (upper panels) and pulse wave flow doppler (lower panels) in wildtype (WT/WT) and heterozygote *Gabarapl1*-KO mice (KO/WT). **f**, Systolic function was maintained in heterozygote *Gabarapl1*-KO mice (WT: n=8, KO/WT n=11-12 animals/group, unpaired 2-sided T-test). **g**, Heterozygote *Gabarapl1*-KO mice exhibit diastolic dysfunction, shown by increased E/e’ ratio (WT: n=8, KO/WT n=12 animals/group, unpaired 2-sided T-test, p=0.043). **h**, Cardiac glycogen content is increased in neonate (p2, age 2 day, WT: n=12, KO/WT: n=9 animals/group, p=0.034) and adult (age 20 week) heterozygote *Gabarapl1*-KO mice (WT: n=7, KO/WT: n=5 animals/group, unpaired 2-sided T-test, p=0.004). **i**, Periodic-acid Schiff (PAS) stained myocardial sections of 20 week old heterozygote *Gabarapl1*-KO mice show increased glycogen (pink-purple) staining (scale bar 50µm). Representative micrographs were selected from a gallery of images acquired from 2 animals/group. Data presented as mean ± s.e.m. *p<0.05. See also Extended Data Figure 2 and Extended Data Table 3.

Other data indicate that in general, while autophagy receptor proteins are degraded in the phagolysosome, ATG8 proteins are at least partially recycled to the cytosol to supply ongoing phagosome formation.^25,26^ Thus, we reasoned that cardiac-directed gene therapy to provide augmented cytosolic GABARAPL1 production could remediate cardiomyocyte glycogen flux dysregulation in diabetes and exert a beneficial effect on diastolic dysfunction.

### AAV Atg8 gene delivery rescues cardiomyocyte and cardiac diastolic dysfunction in diabetic mice

To determine whether cardiac-specific *Gabarapl1* gene delivery could mitigate progression of glycogen accumulation and diastolic dysfunction in diabetes, an AAV9-cTnT-Gabarapl1 (AAV-Gab) viral gene therapy approach was employed (Fig. 4a). Initially we tested if *Gabarapl1* overexpression could modulate glycogen accumulation in cardiomyocytes *in vitro*. Cultured NRVMs were transduced with AAV9-cTnT-Gabarapl1 to induce *Gabarapl1* overexpression (∼1.8 fold increase, Extended Data Figure 3a). Glycogen accumulation in NRVMs induced by exposure to media containing high glucose was suppressed by AAV-Gab gene delivery (Fig. 4b, Extended Data Table 5), confirming Gabarapl1 as a viable target for ‘rescuing’ glycogen dysregulation in a simulated diabetic setting, albeit in an NRVM model known to have high glycolytic dependency.^27^ Next, we investigated Gabarapl1 gene delivery in T2D mice *in vivo*. AAV-Gabarapl1 construct schematic and validation of *in vivo* overexpression are presented in Fig. 4a and Extended Data Fig. 3b respectively. AAV-Gab virus titres were injected by tail vein into T2D and control mice after 14 weeks of high fat high sugar dietary treatment (Extended Data Fig. 3c). Mice were monitored for an additional treatment period of 12 weeks. As expected, the cardiac-specific AAV-Gab virus had no effect on systemic measures of body weight, glucose tolerance and hyperglycemia (Fig. 4c,d, Extended Data Table 6). In contrast, *Gabarapl1* gene delivery had a major cardiac effect. T2D-associated cardiac glycogen accumulation was reversed and diastolic dysfunction (i.e. E/e’) was normalized in T2D mice injected with AAV-Gab at 12 weeks post-treatment (Fig. 4e,f and Extended Data Fig. 3d). E/e’ was negatively correlated with GABARAPL1 protein expression in the membrane-enriched fraction of cardiac protein homogenate (Fig. 4g). Systolic function was unaffected by diabetes and AAV9 treatment (Extended Data Fig. 3d and Extended Data Table 6). Gabarapl1 gene delivery decreased AMPK signaling (phosphorylated (Thr172) to total AMPK ratio) and myocardial lipid deposition (oil red O staining) in T2D mice (Extended Data Fig. 3e-f). Thus cardiac-targeted *Gabarapl1* gene delivery dramatically suppressed myocardial glycogen accumulation and restored diastolic dysfunction in a dietary model of type 2 diabetes progression.

**Fig. 4.**
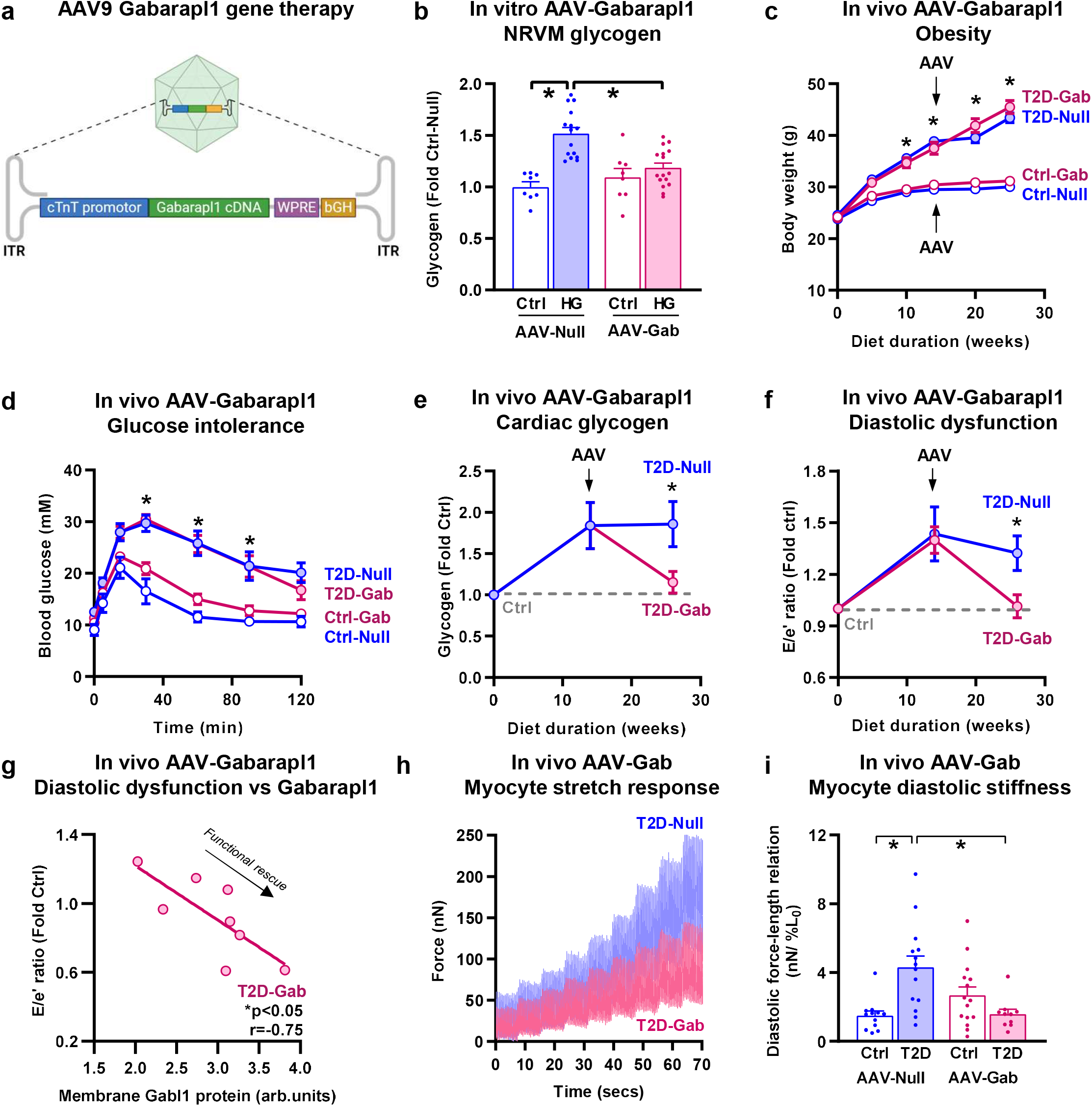
AAV Atg8 gene delivery rescues cardiomyocyte and cardiac diastolic dysfunction in diabetic mice. **a**, Schematic of AAV9 cardiac Gabarapl1 gene delivery construct. Made with Biorender. **b**, Cardiomyocyte glycogen is increased with exposure to high glucose in Null-treated but not AAV-Gabarapl1 treated cells (Ctrl-Null: n=8, HG-Null: n=15, Ctrl-Gab n=8, HG-Gab: n=16 independent culture wells/group, 2-way ANOVA with Bonferroni post-hoc, Null Ctrl v HG p<0.0001, HG Null vs Gab p=0.0001). **c**, In vivo cardiac-specific AAV9-cTnT-Gabarapl1 gene delivery does not affect the development of obesity in T2D mice (induced by high fat high sugar diet; open circles control diet, shaded circles T2D; Ctrl-Null, Ctrl-Gab, T2D-Null: n=10, T2D-Gab n=14 animals/group, repeated measures ANOVA with Bonferroni post-hoc, p<0.0001). **d**, T2D-induced glucose intolerance is not affected by in vivo cardiac-specific AAV-Gab gene delivery (open circles control diet, shaded circles T2D; Ctrl-Null n=5, Ctrl-Gab, T2D-Null, T2D-Gab: n=6 animals/group, repeated measures ANOVA with Bonferroni post-hoc, p<0.01). Baseline (time 0) fasted blood glucose levels are significantly elevated with T2D (2-way ANOVA T2D effect, p=0.019). **e**, Cardiac glycogen accumulation in T2D mice is rescued by cardiac Gabarapl1 gene delivery (13 weeks post-AAV injection, presented as fold change vs Ctrl-fed mice for each virus group, pre-AAV n=10 animals/group, post-AAV T2D-Null n=11, post-AAV T2D-Gab n=13, analyzed by 2-way ANOVA with Bonferroni post-hoc, post-AAV T2D-Gab vs T2D-Null p=0.0276). **f**, T2D-induced diastolic dysfunction (E/e’ ratio, echocardiography) is rescued by cardiac Gabarapl1 gene delivery in AAV-transduced mice (presented as fold change vs Ctrl mice for each virus group. T2D-Null pre-& post-AAV n=10, T2D-Gab pre-AAV n=14, T2D-Gab post-AAV n=13 animals/group, analysed by repeated measures ANOVA, post-AAV T2D-Gab vs T2D-Null p=0.028). **g**, Diastolic function (E/e’) is inversely correlated with Gabarapl1 protein expression in the membrane-enriched fraction of protein homogenates from T2D mice (high fat diet) injected with AAV-Gab virus (12-13 weeks post-AAV injection, r, Pearson correlation coefficient). **h**, Single intact cardiomyocyte force traces (exemplar) show rescued diastolic force (an indicator of passive stiffness) with cardiac Gabarapl1 gene delivery in diabetic mice (AAV9-cTnT-Gabarapl1; T2D, high fat diet; 24 weeks post-AAV injection). **i**, T2D-induced cardiomyocyte diastolic stiffness is rescued by cardiac Gabarapl1 gene delivery in diabetic mice (end-diastolic force-length relation slope; isolated stretched cardiomyocytes from mice at 24 weeks post-AAV injection, high fat diet, Ctrl-Null n=14, T2D-Null n=16, Ctrl-Gab n=15, T2D-Gab n=11 cells/group. Analyzed by 2-way ANOVA with Bonferroni post-hoc, T2D-Null vs Ctrl-Null p=0.022, T2D-Gab v T2D-Null p=0.032). Data are presented as mean ± s.e.m, *p<0.05. See also Extended Data Figure 3 & Extended Data Table 5&6.

To identify possible mechanisms of diastolic functional impairment and remediation we evaluated the performance of single cardiomyocytes obtained from the hearts of AAV-Gab & AAV-null mice after the completion of the treatment period (animal age ≥36 weeks). Recently we have demonstrated *in vitro* that a component of diastolic relaxation pathology in cardiometabolic disease can be attributed to impaired mechanical performance of cardiomyocytes.^28^ Single intact cardiomyocytes were isolated from *ex vivo* mouse hearts, by enzymatic perfusion & dissociation procedures. Each cardiomyocyte was attached to a pair of glass fibres coated with a bio-adhesive – one fibre used to apply stretch steps and the other to a force transducer. Single cardiomyocyte nano-mechanical properties were measured using a protocol of calibrated stretch steps applied during pacing (Extended Data Fig. 3g).^28^ Records superimposed from 2 representative cardiomyocytes from each of the AAV-treated groups shows that with each stretch step both the diastolic and systolic force increases (Fig. 4h). The diastolic force-length relation for each myocyte was calculated from the gradient of the line fitted to the data points collected prior to transition to a new step (normalized to % stretch relative to initial length, L_0_, Fig. 4h,i). The mean data show that the slope of the end-diastolic force-length relation (nN/%L_0_: an index of myocyte ‘stiffness’) was significantly increased in cardiomyocytes from T2D hearts (more than doubled), and with AAV-Gab treatment the mean stiffness gradient in T2D cardiomyocytes was not different to control (Fig. 4i). These data provide a link between cardiomyocyte stiffness and glycophagy disturbance in diabetic heart disease.

Next, we evaluated cardiomyocyte Ca^2+^ handling in T2D hearts treated with AAV-Gab. Ca^2+^ transient amplitude and time to peak were not affected by T2D or AAV-Gab treatment (Fig. 5a,b). T2D mice treated with AAV-Gab exhibited faster Ca^2+^ transient decay relative to null-treated T2D mice (Fig. 5c,d), an effect which was particularly pronounced at the later stage of transient decay (90% to baseline). These findings suggest that glycophagy intervention in T2D mice is linked to accelerated Ca^2+^ transient decay.

**Fig. 5.**
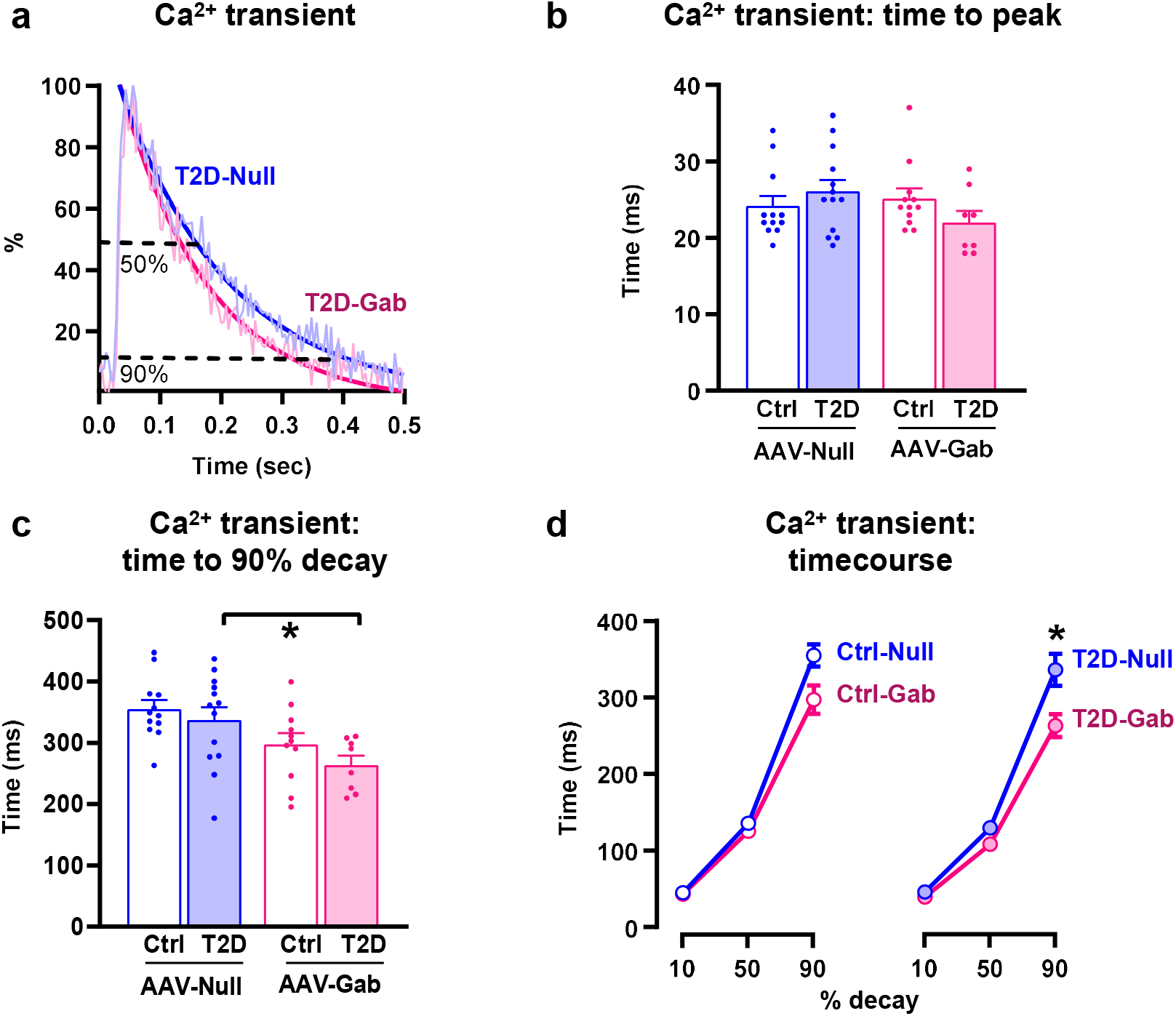
AAV Atg8 gene delivery *in vivo* expedites cytosolic Ca^2+^ removal during cardiomyocyte relaxation in diabetic mice. **a**, Exemplar Ca^2+^ transients recorded at basal length from AAV-Null or AAV-Gab T2D mice 24 weeks post-injection expressed as a percentage of peak amplitude (avg 10 contraction cycles, 2Hz). A mono-exponential decay fit is depicted with dark bold solid lines for each transient and 50 and 90% from peak to baseline are indicated with black broken lines. **b**, Cardiomyocyte Ca^2+^ time to peak is unchanged by T2D or Gabarapl1 gene delivery (Ctrl-Null n=12, T2D-Null n=13, Ctrl-Gab n=12, T2D-Gab n=8 cells, analyzed by 2-way ANOVA). **c**, Cardiomyocyte Ca^2+^ transient 90% decay is faster in diabetic mice treated with Gabarapl1 gene delivery (Ctrl-Null n=12, T2D-Null n=13, Ctrl-Gab n=11, T2D-Gab n=8 cells, analyzed by 2-way ANOVA with Bonferroni post-hoc, T2D-Gab v T2D-Null p=0.021) **d**, Accelerated cytosolic Ca^2+^ removal becomes more pronounced at the later stage of Ca^2+^ transient decay. Note, 0% decay is time to peak. (Ctrl-Null n=12-13, T2D-Null n=13-14, Ctrl-Gab n=11-13, T2D-Gab n=8-9 cells, analyzed by 2-way ANOVA with Bonferroni post-hoc, T2D-Gab v T2D-Null 90% decay p=0.031). Data are presented as mean ± s.e.m, *p>0.05. See also Extended Data Figure 3 & Extended Data Table 6.

### AAV Atg8 gene delivery improves diastolic performance in ‘diabetic’ human iPSC-derived cardiac organoids

To explore the translational potential of *Gabarapl1* gene delivery in improving diastolic dysfunction in diabetic patients, the *Gabarapl1* viral gene delivery approach was applied to human cardiac organoids in an *in vitro* setting of simulated diabetes (‘hyperglycemia’). Three-dimensional engineered heart muscle tissues were formed from human pluripotent stem cells.^26^ The hCO were grown in wells around two semi-flexible posts to support longitudinal tissue growth and stimulated to contract. Calibrated post movement within each well was tracked to measure the force developed by the contracting hCO tissues (in µN).^29,30^ Human iPSCs have higher baseline relative glycogen content than intact hearts, which may reflect differences in metabolic substrate preferences. Confocal microscopy shows α-actinin-stained contractile tissue is evident throughout the cardiac organoid structure (Fig. 6a). With transmission electron microscopy increased glycogen deposits (black particulates throughout the cytosol) were apparent in organoid tissues cultured in high glucose conditions along with evidence of glycophagosome membranous formations (Fig. 6b). High glucose-induced glycogen accumulation was confirmed in the precursor human pluripotent stem cells (Extended Data Fig. 3h). In human cardiac organoids transduced with AAV-Gab, high glucose incubation conditions were linked with delayed force relaxation (diastolic dysfunction), which was normalized with AAV-Gab gene delivery (Fig. 6c,d). In these cardiac organoids, systolic function was preserved under all conditions (Fig. 6e). These data confirm that *Gabarapl1* gene delivery attenuates diastolic dysfunction in human cardiac organoids, thus strengthening the case for translational potential.

**Fig. 6.**
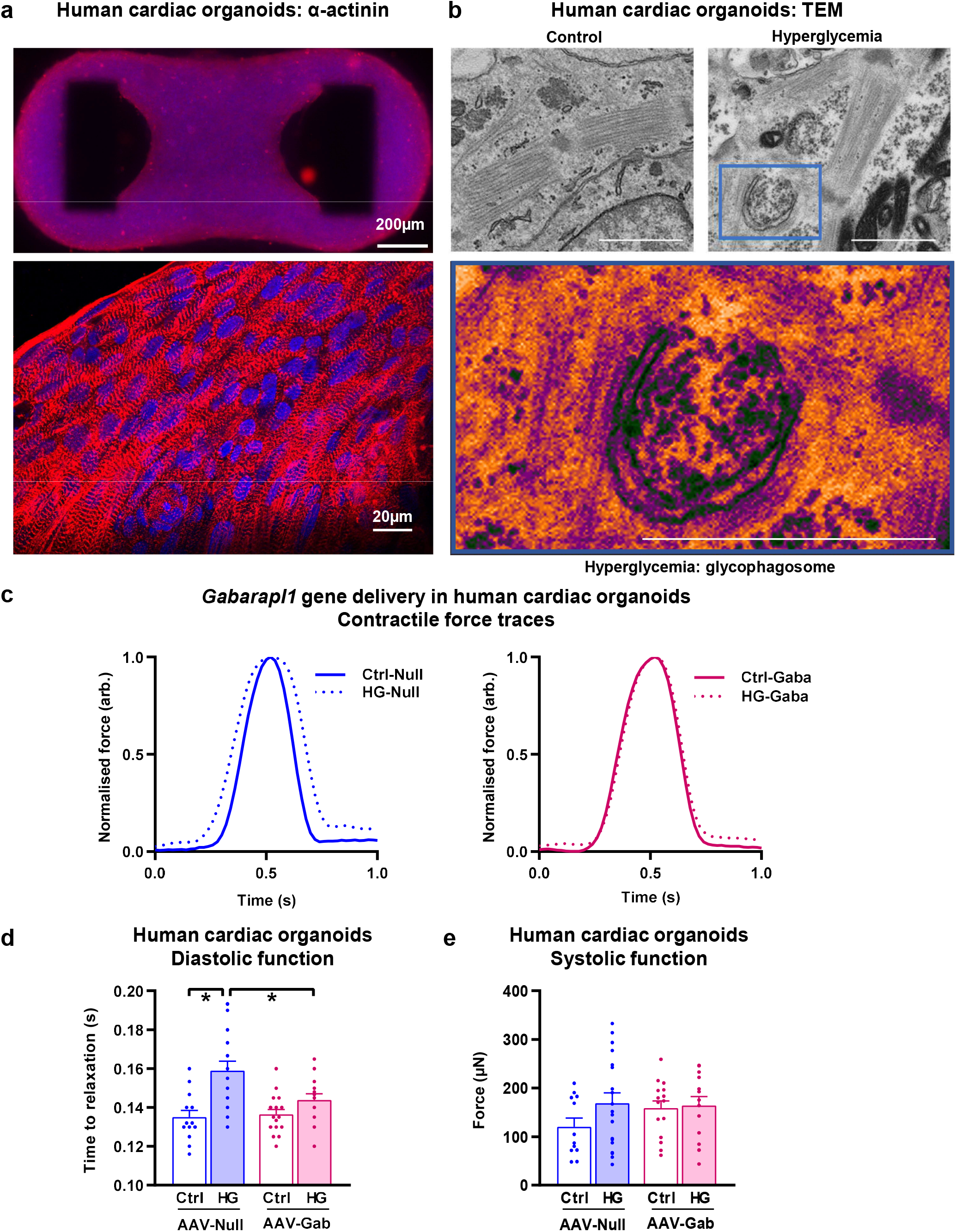
AAV Atg8 gene delivery improves diastolic performance in ‘diabetic’ human iPSC-derived cardiac organoids. **a**, Illustrative confocal image of a human pluripotent stem cell-derived cardiac organoid, stained for α-actinin (red) and DNA (blue, Hoescht stain). **b**, Transmission electron microscopy images of cardiac organoids showing increased glycogen deposits (black dots) in HG-treated organoids. High magnification image of a glycophagosome in an HG-treated organoid. Scale bar, 1μm. Representative electron micrographs were selected from a gallery of 4 images for control and 5 images for hyperglycemia conditions. **c**, Exemplar force traces from human cardiac organoids, normalized systolic force. **d**, HG-induced diastolic dysfunction (delayed time to 50% relaxation of force) is rescued by AAV9-Gabarapl1 transduction in human cardiac organoids (Ctrl-Null n=13, HG-Null n=19, Ctrl-Gab n=16, HG-Gab n=14 organoids, analyzed by 2-way ANOVA with Bonferroni post-hoc, HG-Null v Ctrl-Null p<0.0001, HG-Gab v HG-Null p=0.012). **e**, Systolic function (active force production) is unchanged with HG and Gabarapl1 gene delivery (Ctrl-Null n=12, HG-Null n=19, Ctrl-Gab n=15, HG-Gab n=13 organoids, analyzed by 2-way ANOVA). Data are presented as mean ± s.e.m, *p<0.05. See also Extended Data Figure 3.

## DISCUSSION

In summary, these findings provide compelling evidence that cardiomyocyte specific regulation of glycophagy constitutes a major myocardial metabolic axis of pathophysiologic importance. Here, using a dietary model of type 2 diabetes exhibiting diastolic dysfunction, we show that cardiac-targeted gene therapy delivering the ATG8 family GABARAPL1 partner of STBD1 (the receptor which binds the glycogen cargo for glycophagosome capture) resolves cardiomyocyte glycogen accumulation and alleviates diastolic dysfunction.

The mechanisms linking cardiac glycophagy intervention to improved diastolic function in T2D are likely multifaceted. Our recent biomechanical experimental work has demonstrated that diastolic dysfunction measured in vivo has a substantial origin in cardiomyocyte ‘stiffness’ (relaxation deficit).^28^ In this study, our findings indicate that the Gabarapl1-mediated glycophagy intervention accelerated cytosolic Ca^2+^ clearance and normalized cardiomyocyte diastolic stiffness in T2D mice. Evidence of glycolytic dependence of key molecular Ca^2+^ transporters involved in cardiomyocyte diastolic relaxation has been reported, including the sarcoplasmic reticulum Ca^2+^-ATPase, SERCA2.^12,13^ We speculate that cardiac Gabarapl1 gene therapy promotes glycophagosome delivery of glycolysis substrate ‘packages’ (i.e. via optimal sarcomeric positioning of lysosomal glycogenolysis) to enhance SERCA2-mediated Ca^2+^ reuptake and cardiomyocyte relaxation. Our finding that T2D mice treated with Gabarapl1 exhibit reduced glycogen accumulation and lower AMPK activity is consistent with the contention that targeted upregulation of glycophagy augments glycogenolysis and promotes ATP availability. Further, decreased AMPK activity with Gabarapl1 gene therapy is associated with relieved burden of accumulated cytosolic lipids in the T2D model. Cardiac lipotoxicity is considered a ‘hallmark’ of the diabetic phenotype and has been linked to diastolic dysfunction^31^ – although the mechanisms are not well understood. A graphic to illustrate the potential role of glycophagy in cardiomyocyte functional support has been included in Figure 7.

**Fig. 7.**
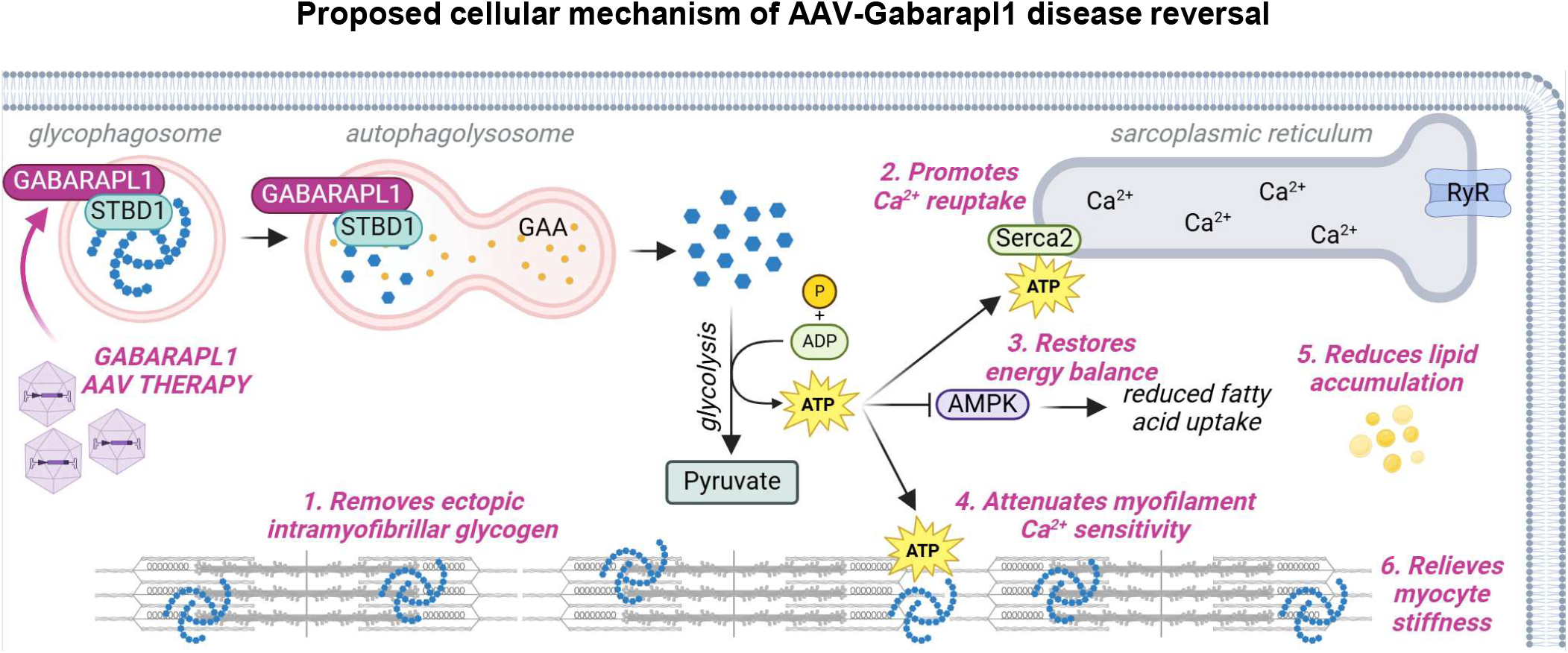
Proposed cellular mechanism. A hypothetical mechanism of glycophagy role in physiology and pathology involvement in diabetic heart disease. Created with Biorender.com.

Recently glycophagy has emerged as a selective autophagic route operating in parallel with other autophagy types including macro/aggrephagy (protein aggregates), mitophagy (organelle turnover) and others.^3^ It is likely that autophagy pathways involve some ATG8 redundancy and that the cargo capture may be selective but not exclusive. While structural analyses have demonstrated that the STBD1-GABARAPL1 receptor-ATG8 partner pair have the highest binding affinity, other GABARAP-subfamily proteins also exhibit STBD1 binding (albeit at lower affinity^19^). Such redundancy would confer useful metabolic flexibility – a feature of particular importance in the heart.

It is possible that STBD1 binding affinities with glycogen and GABARAPL1 differ in disease contexts constituting an important factor influencing glycophagy activity. Immunolabelling to determine GABARAPL1 interactions is limited by poor antibody specificity,^14,32^ and protein tagging is required for accurate visualization. *In vitro* fluorescence polarization assays with FITC-labeled glucose strands revealed that STBD1 preferentially binds to linear glucose polymers with longer glucose units.^19^ The concept that glycophagy favors less-branched glycogen polymers has been advanced in the literature,^4,19^ but whether this is a factor determining glycophagy fate in settings of cardiometabolic stress has not been investigated. Advanced biochemical knowledge of STBD1 has been mostly derived from non-cardiac settings. An understanding of the interplay between STBD1-glycogen binding and the dynamic glycolysis demands of the cardiac cycle would be informative.

Disturbed glycophagy is unlikely to be a condition unique to diabetic heart disease, and may be a factor in a range of conditions where diastolic dysfunction is prominent, including familial hypertrophic cardiomyopathies and heart failure with preserved ejection fraction (HFpEF^33^). Systematic studies are required to assess glycophagy status in other cardiac disease phenotypes.

### Future Directions & Limitations

We have previously identified that fasting-induced glycogen accumulation and glycophagy regulation is accentuated in female hearts,^34^ and early studies showed that the Gabarapl1 gene has an estrogen response element.^35^ Thus glycophagy targeting may be even more efficacious in females and sex/gender-specific studies of glycophagy intervention will be informative. In the present study, the proposition that glycophagy, glycogen and diastolic function are closely linked in cardiac disease was supported by in vitro studies that involved female origin cells (NRVMs, iPSC-derived organoids). Sex/gender comparative studies of cardiac efficacy of gene therapy are important to pursue in the future. In this study AAV Gabarapl1 gene delivery was financially feasible and targetable for mice. Clinical trials investigating the long-term safety and efficacy of gene delivery are underway and will be informative for future human translation for glycophagy gene therapy.^36^ Our finding that *Gabarapl1* deficiency is evident in cardiac tissue from patients with diabetes provides a first step towards validating the mechanism in a clinical setting. Further work is required to fully characterize glycophagy processes in the human heart and translate these findings to the clinic.

### Conclusions

Almost three decades since Ohsumi’s seminal autophagy discovery work^37^, to our knowledge, an autophagy-specific therapy has yet to be clinically tested.^38^ As autophagy disturbance has been implicated in a broad range of disease conditions (affecting both cell survival and death), a major challenge is to devise a precision approach which avoids generalized negative health impacts. Our study provides the pre-clinical basis for demonstration of cardiac-selective glycophagy as a viable treatment target in cardiac metabolic pathology and offers potential for development of interventional therapeutic strategies in diabetic heart disease.

## METHODS

### Animals

Animal experiments were performed at the University of Auckland, the University of Melbourne, and Cedars-Sinai Medical Center. All animal experiments were approved by the University of Auckland Animal Ethics Committee or Cedars-Sinai Animal Ethics Committee and complied with the guidelines and regulations of the Code of Practice for the Care and Use of Animals for Scientific Purposes. Age-matched male animals were randomly assigned to experimental groups and were housed ≥ 2 animals per cage in a temperature-controlled environment 21-23 ^°^C with 12 hour light/dark cycles. At the start of all experiments, all animals were drug/test naïve. Unless otherwise specified, animals were fed regular chow. At study completion non-fasted animals were euthanized and tissues were collected between 10am and 1pm.

#### Acute metabolic stress rats

Metabolic stress was induced in male Sprague Dawley rats (sourced from Charles River) at 8 weeks of age via a single 55mg/kg tail vein injection of streptozotocin (STZ; Sigma) and monitored for 2 days, animals were not insulin-treated. The acute STZ-treatment in the rat provides a setting of rapid onset hyperglycemic metabolic challenge for interrogation of early glycogen disturbance at 48hr post treatment (not evident in STZ mice). Non-fasted blood glucose was determined by tail prick blood collection analysed by accu-chek glucometer.

#### Type 2 diabetic high fat high sugar diet mice

Obesity and type 2 diabetes was induced in male C57Bl/6J mice (sourced from the University of Auckland breeding facility) by a high fat high sugar dietary intervention commencing at 8-9 weeks of age.^39^ After a 1 week transitional feeding period, animals were fed a high fat diet with high sugar (43% kcal from fat, 200g/kg sucrose, SF04-001, Specialty Feeds) or control reference diet (16% kcal from fat, 100g/kg sucrose, custom mouse AIN93G control diet, Specialty Feeds) for 14-15 weeks. High fat high sugar-fed mice exhibit obesity, mild hyperglycemia and glucose intolerance (Extended Data Table 2). This mouse model recapitulates the clinical phenotype of diabetic heart disease and provides a suitable setting for AAV intervention.

#### Global Gabarapl1-KO mice

CRISPR-Cas9 Gabarapl1-knockout mice were generated by the Melbourne Advanced Genome Editing Centre on a C57Bl/6J background using gRNAs specified in Extended Data Table 3. Next generation sequencing confirmed excision of exons 2-4 from the *Gabarapl1* gene in founder mice. Mice used in these studies were male and selected from 8 independent founder lines for each knockout model (Extended Data Fig. 2). Mice were 10 weeks old unless otherwise specified in the Figure legend.

#### Type 2 diabetic high fat high sugar diet mice with AAV-Gabarapl1 gene delivery

To evaluate the effect of cardiac-specific *Gabarapl1* gene delivery in diabetic heart disease, high fat high sugar- and control-fed male C57Bl/6J mice were randomized to treatment group after 14 weeks of the dietary intervention (at 22 weeks old) and received a single tail vein injection of 10^12^ gc/mouse AAV9-cTnT-*Gabarapl1* or AAV9-cTnT-Null virus (Vector Biolabs). The mice continued the high fat high sugar or control diet for a further 14 weeks (glycogen analysis) or 28 weeks (cardiomyocyte stiffness analysis).

### Human myocardial samples

Right atrial appendage samples were obtained from non-diabetic and type 2 diabetic patients (aged 50– 80 years, male and female) at the time of elective cardiac surgery either at the Royal Melbourne Hospital, Australia, or Dunedin Hospital, New Zealand (patient characteristics are summarized in Extended Data Table 4). Collection of clinical samples was approved by the Melbourne Health Human Research and Ethics Committee and the Human and Disability Ethics Committee of New Zealand (LRS/12/01/001), and all patients provided informed consent.

### Human pluripotent stem cell-derived cardiac organoids

Human pluripotent stem cells were maintained and differentiated into cardiomyocytes (hPSC-CMs)^29,30^ with ethical approval from The University of Queensland’s Medical Research Ethics Committee under the regulations of the National Health and Medical Research Council of Australia (NHMRC). iPSCs were originally obtained from ATCC (#BXS0116) and were derived from bone marrow CD34+ cells obtained from a healthy, 31-year-old, White, female donor. hPSC-CMs were dissociated and seeded onto gelatin coated coverslips at 100,000 cells/cm^2^ in α-MEM GlutaMAX (Thermo Fisher Scientific), 10 % Fetal bovine serum (FBS; Thermo Fisher Scientific), 200 μM L-ascorbic acid 2-phosphate sesquimagnesium salt hydrate (Sigma) and 1 % penicillin/streptomycin (Thermo Fisher Scientific). Human cardiac organoids (hCOs) were generated from hPSC-CMs using Heart-Dyno culture inserts and cultured in maturation media.^29^ Briefly, cells were allowed to condense around two separated semi-flexible posts forming hCOs (∼1mm length). The hCO tissues grown in each well were stimulated to contract, pulling against the posts. The resistance of the posts to movement during contraction was calibrated based on prior force transducer validation. Calibrated post movement within each well was tracked to measure the force developed by the each contracting hCO tissue (in µN).^29,30^ To ensure that culture conditions were robust and applicable to multiple hPSC lines, we optimized the serum-free medium and matrix composition. Important protocol standardization measures included removal of TGFb-1, increased collagen content with Matrigel, and increased culture time to 5 days to allow for organoid tissue condensation. Robust and functionally reproducible organoids were derived from all hPSC lines tested. Additional papers further characterizing the human-iPSC-derived organoids and functional evaluation are available.^29,30^

To obtain images of contractile protein ultrastructure, organoids were immunostained with alpha-actinin and hoescht blue (DNA stain) and imaged with a confocal microscope. At Day 9 hCOs were fully formed and ‘matured’ and were treated with AAV9-cTnT-*Gabarapl1* or AAV9-cTnT-Null virus (5×10^9^ gc/hCO; Vector Biolabs) for 72 hours in maturation media. hCOs were subsequently washed and cultured in control (Ctrl: maturation media with 1 mM glucose with 24 mM mannitol) or simulated hyperglycemic (HG: maturation media with 25 mM glucose) conditions for 4 days to allow time for glycogen accumulation. At the completion of contraction analysis a subset of cardiac organoids were fixed for electron microscopy.^29^

### Neonatal rat ventricular myocyte culture

Cardiomyocytes from male and female postnatal day 1-2 Sprague Dawley rat (sourced from the University of Auckland breeding facility) hearts were isolated and cultured as previously described.^10^ Briefly, ventricular myocytes were isolated by enzymatic dissociation with trypsin and collagenase. Isolated cells were centrifuged (10 minutes, 1000 g), and resuspended in minimum essential media (MEM, Life Technologies) supplemented with 10 % newborn calf serum (NCBS), essential and non-essential amino acids, antibiotic-antimycotic reagent, 2 mM L-glutamine, and MEM vitamins (Life Technologies) and 26 mM NaHCO_3_ and 98 µM bromo-deoxyuridine (Sigma), and pre-plated for 1.5 hours at 37 °C to obtain a cardiomyocyte-rich fraction. Cells were plated at a cell density of 1250 cells/mm^2^ and incubated at 37 ^°^C with 5 % CO_2_. To evaluate the effect of specific gene knockdown, NVRMs were switched to antibiotic-free MEM-10%NBCS media with 100nM siRNA (rat siGabarapl1; siGENOME SMARTpool, Dharmacon, Extended Data Table 3) vs non-targeting scrambled siRNA with Lipofectamine RNAiMAX (Invitrogen) for 16 hours. Following transfection, NRVMs were cultured for 24 hours in MEM-10%NBCS with antibiotics prior to 24 hours serum-free Dulbecco’s modified essential media (DMEM; Sigma) with 5mM glucose and 1nM insulin). Cells were lysed with radioimmunoprecipitation buffer (RIPA, ThermoFisher) for molecular analysis. For *Gabarapl1* overexpression studies, following isolation NRVMs were cultured in MEM-NBCS as described above, for 24 hours, then switched to antibiotic-free, serum-free Opti-MEM (Thermo Fisher Scientific) with AAV9-cTnT-*Gabarapl1* or AAV9-cTnT-Null virus (10,000 gc/cell; Vector Biolabs) for 24 hours. Following transduction, NRVMs were cultured for 48 hours in standard DMEM (with supplements as described above, and 5 mM glucose) prior to 24 hours experimental conditions of control glucose (Ctrl: 5 mM glucose, 25 mM mannitol) vs high glucose (HG: 30 mM glucose). At the completion of the experimental period, cells were lysed with RIPA buffer (Thermo Fisher Scientific) for molecular analysis.

### Glucose tolerance testing

Glucose tolerance testing was performed in T2D-AAV mice following 6 hours fasting.^40^ Baseline blood glucose levels were measured using an Accu-Chek glucometer with a small blood sample obtained from a needle prick to the tail vein. Glucose (1.5 g/kg body weight) was injected i.p. and blood glucose was measured at 5, 15, 30, 60 and 90 minutes after the glucose injection.

### Echocardiography

Animals were anaesthetized with isoflurane and transthoracic echocardiography was performed using the GE Vivid 9 Dimension echocardiography platform with a 15 MHz i13L linear array transducer (GE Healthcare).^39^ Left ventricular (LV) m-mode two-dimensional echocardiography was performed in parasternal short axis view to measure LV wall and chamber dimensions to derive systolic function parameters: % ejection fraction ((end diastolic volume – end systolic volume)/end diastolic volume) ×100), and % fractional shortening ((LV end diastolic diameter – LV end systolic diameter)/(LV end diastolic diameter) ×100). Pulse wave Doppler and tissue Doppler imaging were acquired from the apical 4 chamber view to assess LV diastolic function parameters: velocity of early mitral inflow (E) and early diastolic velocity of mitral annulus (e′) and E/e′ ratio. Three consecutive cardiac cycles were sampled for each measurement taken, and analysis was performed in a blinded manner.

### Mouse cardiomyocyte isolation, Ca^2+^ handling and nano-mechanical analysis

Adult mouse cardiomyocytes were isolated,^41^ and assessed for myocyte stiffness properties and Ca^2+^ handling. Cardiomyocytes were loaded with Fura2-AM (1 µM) and attached to glass rods (Myotak, Ionoptix) and lifted from the coverslip using motorized micromanipulators (MC1000E, Siskiyou Corporation). Cardiomyocytes were paced (2 Hz, 2.0 mM Ca^2+^, 37 °C) and subjected to progressive longitudinal stretch (Nano-drive, Mad City Labs). Sarcomere length/shortening and force development were measured (force transducer stiffness 20 N/m; Myostretcher, Ionoptix) in parallel with acquisition of Ca^2+^ transients (Fura2 microfluorimetry F360:380nm ratio; IonOptix). Diastolic force-length relationships were constructed by plotting diastolic force against % cardiomyocyte stretch. All indices were analyzed off-line using IonWizard software (IonOptix) and were determined after averaging 10 steady-state transients for each myocyte.

### Unloaded adult rat cardiomyocyte Ca^2+^ handling

Adult rat cardiomyocytes from STZ (8 weeks post-STZ injection) and control rats were isolated for Ca^2+^ analysis.^28^ Cardiomyocytes were superfused with a HEPES-Krebs buffer (in mM: 146.2 NaCl, 4.69 KCl, 0.35 NaH_2_PO_4_H2O, 1.05 MgSO_4_7H_2_O, 10 HEPES, 2 CaCl_2_) under physiological conditions (4Hz field-stimulation, 37°C) and analyzed for Ca^2+^ handling (Fura2, 1μM, microfluorimetry F360:380nm ratio; IonOptix). All indices were analyzed off-line using IonWizard (IonOptix) and were determined after averaging 10 steady-state transients for each myocyte.

### Molecular analyses

#### Glycogen content

Glycogen content was measured by digesting an aliquot of tissue homogenate or cell lysate with amyloglucosidase (#10102857001, Roche) at 50 °C for 60 minutes in 1 % triton-X, 0.1 M Na acetate, pH 6.0. Following centrifugation at 16,000 g, 4 °C for 2 minutes, the supernatant was assayed for glucose concentration using glucose oxidase/peroxidase colorimetric glucose assay with 45 μg/ml o-dianisdine dihydrochloride (Sigma-Aldrich), and absorbance measured at 450 nm. Another tissue homogenate aliquot was processed in parallel without amyloglucosidase to determine background glucose content. Glycogen levels are presented as glucose units (nmol), normalized to protein (mg, determined by Lowry assay) and depicted as relative levels for comparative purposes.

#### Immunoblotting

Sample protein concentrations were determined using a Lowry assay. Prior to immunoblotting, tissue homogenates were prepared in loading buffer (50 mM Tris-HCl (pH 6.8), 2 % sodium dodecyl sulfate, 10 % glycerol, 0.1 % bromophenol blue and 2.5 % 2-mercaptoethanol), and equal amounts of protein were loaded into the SDS-PAGE gel. Antibodies for the following proteins were used: phosphorylated (Ser641) glycogen synthase (ab81230, Abcam), glycogen synthase (3893, Cell Signaling), phosphorylated (Ser14) glycogen phosphorylase (gift from Dr David Stapleton), glycogen phosphorylase (gift from Dr David Stapleton), GABARAPL1 (26632, Cell Signaling), STBD1 (gifted by Dr David Stapleton), phosphorylated (Thr172) AMPK (2535, Cell Signaling) and AMPK (2532, Cell Signaling). Membranes were incubated with anti-rabbit horseradish peroxidase-conjugated secondary antibody (GE Healthcare). The ECL Prime (Amersham, GE Healthcare) chemiluminescent signal was visualized with a ChemiDoc-XRS Imaging device and band intensity quantified using ImageLab software (Bio-Rad). Equal protein loading was confirmed by Coomassie staining of polyvinylidene difluoride membranes (Coomassie Brilliant Blue R-250, Bio-Rad).

#### Quantitative RT-PCR

RNA was extracted from frozen cardiac tissues using the TRIzol® reagent in conjunction with the PureLink™ Micro-to-Midi Total RNA Purification kit (Life Technologies) with on-column DNase treatment (PureLink DNase; Life Technologies). RNA was reverse transcribed as per the manufacturer’s instructions (#18080-051; Life Technologies). Real-time PCR was performed using Sybr-Green (Life Technologies) with primer pairs detailed in Extended Data Table 3. The comparative ΔΔCt method was used to analyze the genes of interest relative to housekeeper gene 18S rRNA. Gene expression was evaluated in RNA extracted from ventricular (animal studies) or atrial (clinical samples) tissue, and the influence of changes in the relative cellular composition of these tissues with disease on transcript levels cannot be excluded.

For assessment of viral copy number in AAV9-treated mouse heart tissue, DNA was extracted as per the manufacturer’s instructions (K0152, Thermo Fisher Scientific). AAV9-treated mouse heart DNA was subjected to digital droplet qPCR as per the manufacturer’s instructions (Biorad) using primers targeting the WPRE component of the viral vector: 5’-CTGGTTGCTGTCTCTTTATGAGGAG-3’ (forward) and 5’-CACTGTGTTTGCTGACGCAACC-3’ (reverse). Viral copies/μg DNA was quantified using QuantaSoft™ Analysis Pro.

#### Glycogen proteomics

Cardiac (left ventricle) and skeletal muscle (quadriceps) tissues were collected from control and acute metabolic stress rats (2 days post-destruction of pancreatic β-cells with 55mg/kg STZ tail vein injection, a time-point when blood glucose levels were >20mM). Briefly, tissues were homogenized in glycogen extraction buffer (in mM: 50 Tris pH 8, 150 NaCl, 2 EDTA) and cellular debris was removed by centrifugation (6000 g, 10 minutes, 4 °C). The supernatant was ultracentrifuged (300,000 g, 60 minutes, 4 °C), and the resulting pellet was resuspended in glycogen extraction buffer and layered over a sucrose gradient (75, 50, 25 % w/v sucrose) and ultracentrifuged (300,000 g, 2 hours, 4 °C). The resulting pellet containing glycogen was resuspended in glycogen extraction buffer.^42^ Glycogen-associated proteins were displaced from the glycogen by addition of α1,4 malto-oligosaccharides (maltodextrin, 50 mg/ml) and ultracentrifuged (400,000 g, 20 minutes, 4 °C). The supernatant containing proteins eluted from the glycogen were reduced (tris(2-carboxyethyl)phosphine (TCEP), 5 mM), alkylated (iodoacetamide, 10 mM), trypsin-digested (1:75 w/w, Promega) and desalted (HLB μElution plate, Waters) in preparation for liquid chromatography tandem mass spectrometry (LC-MS/MS). LC-MS/MS was performed in micro-flow mode (Eksigent 415 LC, 5600+ TripleTOF Sciex) using a trap column (ChromXP C18CL 10×0.3 mm 5 µm 120 Å; flow rate 10 µL/min, 3 minutes), and an analytical column (ChromXP C18CL 150×0.3 mm 3 µm 120 Å; flow rate 5 µL/min, 30 °C). Peptides were eluted in a linear gradient spanning 3-35 % acetonitrile in water (0.1 % Formic Acid) over 60 minutes. MS/MS scans were acquired as previously described.^43^ All raw data from the 5600+ TripleTOF mass spectrometer were converted to mzXML using a Sciex Data converter. Data were analyzed using the Sorcerer 2TM-SEQUEST® algorithm (Sage-N Research, Milpitas CA, USA) searched against the concatenated target/decoy Rat Uniprot 2018 reviewed FASTA database^44,45^ limited to trypsin-digested peptides, 50 PPM parent ion tolerance, 1 Da fragment ion tolerance, carbamidomethyl of cysteine fixed modification and oxidation of methionine variable modification. Quantitation, validation and normalization was performed using Scaffold 4 on proteins defined by a minimum of 2 proteotypic peptides (5 % false discovery rate) with >99.0 % probability (Protein Prophet algorithm^46^). Relative protein quantification was derived from MS/MS data using spectral counting, subjected to normalized spectral abundance factor (NSAF)^47^ and a minimum value of 0 assigned to non-detected proteins. A subset of proteins identified to be i) exclusively detected in control samples, ii) exclusively detected in acute metabolic stress samples and iii) differentially abundant in acute metabolic stress vs control samples (−0.2 ≤ fold change ((MS/control)-1) ≥ 0.2, p<0.05) were used for subsequent analyses. Proteins relating to the contractile apparatus and protein translation GO categories were excluded. Venn diagrams were constructed for cardiac and skeletal muscle datasets to show the number of unique or differentially abundant proteins in control vs acute metabolic stress samples identified from the Scaffold 4 output. Proteins uniquely detected in the acute metabolic stress samples (not detected in control samples) were categorized using Enrichr software^46,48^ into GO categories Carbohydrate Metabolic Processes (GO0005975), Fatty Acid Metabolic Processes (GO0006629) and Amino Acid Metabolic Processes (GO0006520) and Log_2_ protein abundance presented in a Bubble Chart format. GO analysis (Enrichr) of all uniquely detected and differentially abundant proteins identified the top 7 GO Biological Processes categories associated with the glycogen proteome response to acute STZ for cardiac and skeletal muscle tissues. The normalized spectral counts of the uniquely detected and differentially abundant proteins in the GO category Carbohydrate Metabolic Processes were converted to heatmap intensity for graphical representation.

#### STRING functional network analysis

The glycogen-tagging protein, STBD1, identified to be of importance in the glycogen proteome response to acute metabolic stress, was subjected to STRING protein-protein association network analysis V11.0^50^ to identify common pathways or associations for subsequent investigation. Each association identified was based on text-mining and/or experimentally determined evidence in the rat database. The minimum accepted confidence score (minimum probability) for an association was set at 0.7. Primary (max. 10) and secondary (max. 20) interactions were identified and presented as a network graph.

### Histology

Heart mid-sections were fixed in 10% formalin, embedded in optimal cutting temperature (OCT) compound, cryosectioned (10µm) and stained with Oil Red O (30 minutes duration) to identify neutral lipid droplets. The detection of neutral lipid droplets was performed using a brightfield microscope (Zeiss Imager D1, connected to a Zeiss AxioCam MRc5 color camera). The small intense red/brown punctate structures relate to intracellular lipid depots. The dark larger stained structures with less-defined edges may be macro-lipid deposits beyond the plane of focus or may be adherent lipid material from the tissue. The possibility that there may be some staining artefact from debris or residual ORO stain cannot be excluded.

### Statistical analysis and reproducibility

Data are presented as mean ± SEM and statistical analysis was performed using Graphpad Prism V9. A description of biological replicates is provided within the figure legends, measurements were taken from discrete samples. Sample size was selected based on variation observed in previous studies with similar endpoints. Statistical assumptions concerning the data, including normal distribution and similar variation between experimental groups, were examined for appropriateness before statistical tests were performed. Any data points that were identified as statistical outliers (+/-2 SD from the mean) were excluded. All treatments were randomly allocated and investigators were blinded to treatment groups whenever feasible. For comparison between two groups, a 2-sided Student’s T-test was used. For assessment between two independent variables, two-way ANOVA with Bonferroni multiple comparisons post-hoc test was used. For correlation analyses Pearson’s correlation coefficient was used. A p-value of <0.05 was considered statistically significant.

## DATA AVAILABILITY

Source data for all datasets generated are provided. Correspondence and requests for materials should be addressed to L.M.D.D.

## ACKNOWLEDGEMENTS

**General:** We acknowledge D.Stapleton for providing the glycogen phosphorylase antibodies. **Funding:** This work was supported by grants from the National Health and Medical Research Council of Australia (NHMRCA;1027865, 628643, 1082215, 1037320, 1067869, 1157320), the Diabetes Australia Research Trust, Stem Cells Australia, the National Heart Foundation of Australia (NHFA), the New Zealand Marsden Fund (14-UOA-160, 19-UOA-268), the Health Research Council of New Zealand (19/190) and the University of Auckland Faculty Research Development Fund, and the National Institutes of Health, USA (R01 HL155346-01 and R01 HL144509-01). Fellowship support is acknowledged from NHMRCA (JEH, RGP, ERP), and Heart Foundation of Australia (ERP, JEH, KLW). The funders had no role in study design, data collection and analysis, decision to publish or preparation of the manuscript.

## AUTHOR CONTRIBUTIONS

L.M.D.D and K.M.M conceived and designed the experiments. K.M.M, U.V., P.K., C.L.C., J.V.J., L.J.D, G.B.B., A.J.A.R., M.A., X.L., S.L.J., D.J.T., K.R., R.J.M., and R.G.P. performed the experiments. K.M.M, U.V., P.K., C.L.C., J.V.J., L.J.D, G.B.B., A.J.A.R., M.A., X.L., S.J., D.J.T., K.R., K.L.W., R.J.M., J.R.B., E.R.P., and J.E.H. analyzed the data. L.M.D.D, K.M.M., E.R.P., J.E.H., J.E.V., X.H, R-P.X, R.K., T.J.O. and R.A.G. contributed materials/analysis tools. L.M.D.D. and K.M.M wrote the paper.

