## Extended Data Figures + Tables for "Targeted glycophagy ATG8 therapy reverses diabetic heart disease in mice and in human engineered cardiac tissues"

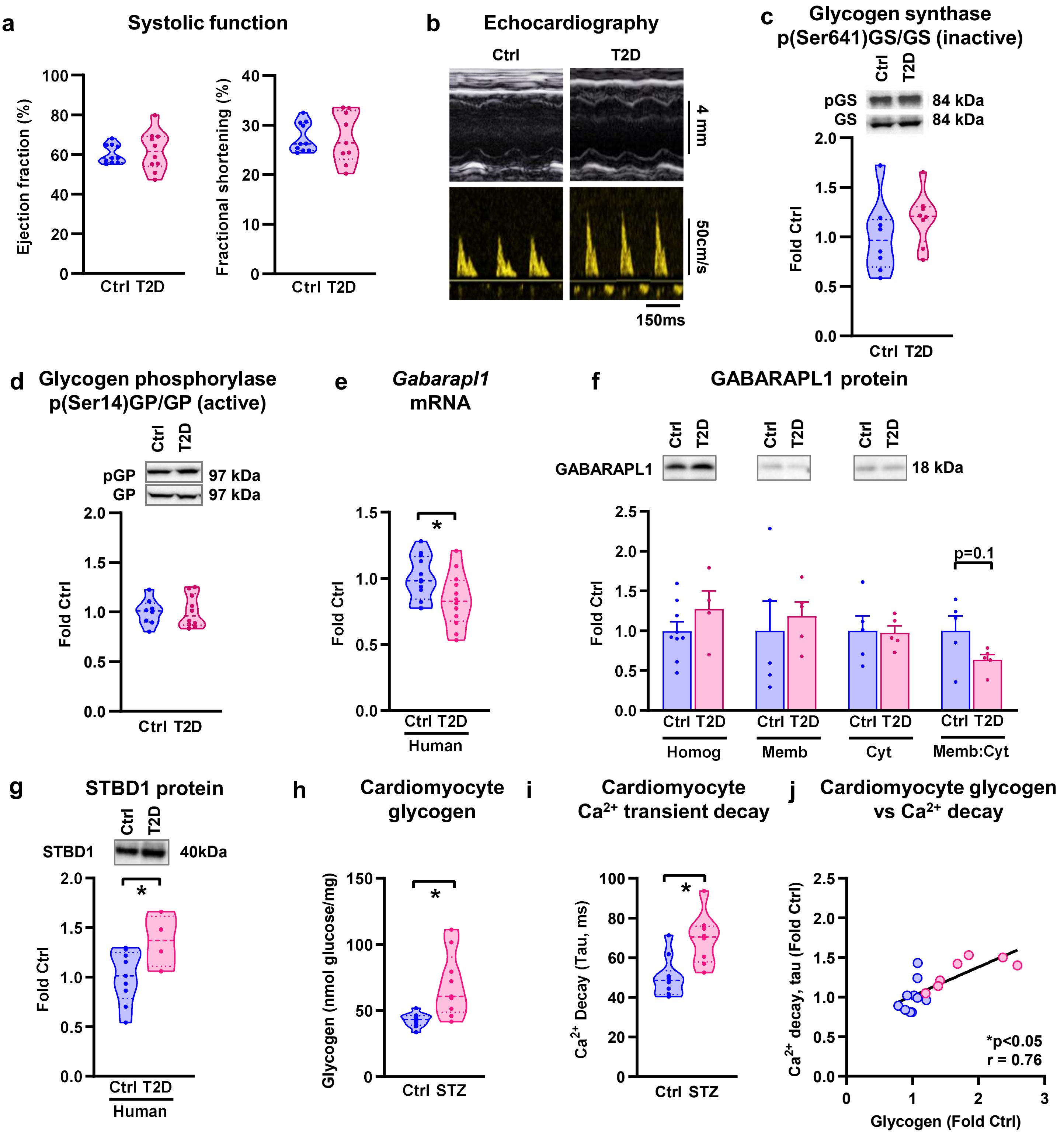


**Extended Data Fig.1 | Cardiac function and glycogen handling in diabetic hearts. a,** Echocardiography M-mode-derived ejection fraction and fractional shortening are unchanged with diabetes in type 2 diabetic (T2D, 14 week high fat sugar diet) vs control (Ctrl) mice (n≥9 animals/group). **b,** Exemplar M-mode and flow Doppler echocardiography traces. **c,** Ratio of phosphorylated glycogen synthase (pGS Ser641) to total glycogen synthase (GS) is unchanged in T2D mouse hearts (n≥8 animals/group, molecular weight indicated in kilodaltons, kDa). **d,** Ratio of phosphorylated glycogen phosphorylase (pGP Ser14) to total glycogen phosphorylase (GP) is unchanged in in T2D mouse hearts (n≥8 animals/group). **e,** Gabarapl1 mRNA is lower in atrial appendage samples from patients with T2D (n=12 patients/group). **f,** GABARAPL1 protein content in fractionated atrial appendage samples from patients with T2D (n≥4 patients/group). **g,** STBD1 protein content is increased in atrial appendage samples from patients with T2D (n≥4 patients/group). **h,** Glycogen content normalized to protein in adult rat cardiomyocyte lysate from Ctrl and streptozotocin (STZ)-induced diabetic rats (8 weeks post-STZ, n≥9 rats/group). **i,** $Ca^{2+}$ time constant of decay (Tau) in isolated adult cardiomyocytes from control and STZ rats (8 weeks post-STZ, n≥8 rats/group). **j,** Correlation of cardiomyocyte glycogen content and $Ca^{2+}$ time constant of decay (Tau) in control (blue) and STZ (pink) rat cardiomyocytes (unloaded cells; r, Pearson correlation coefficient). Data presented as truncated violin plots with median and upper & lower quartiles indicated, or bar graphs mean +/- sem. *p<0.05. Related to Figure 2.

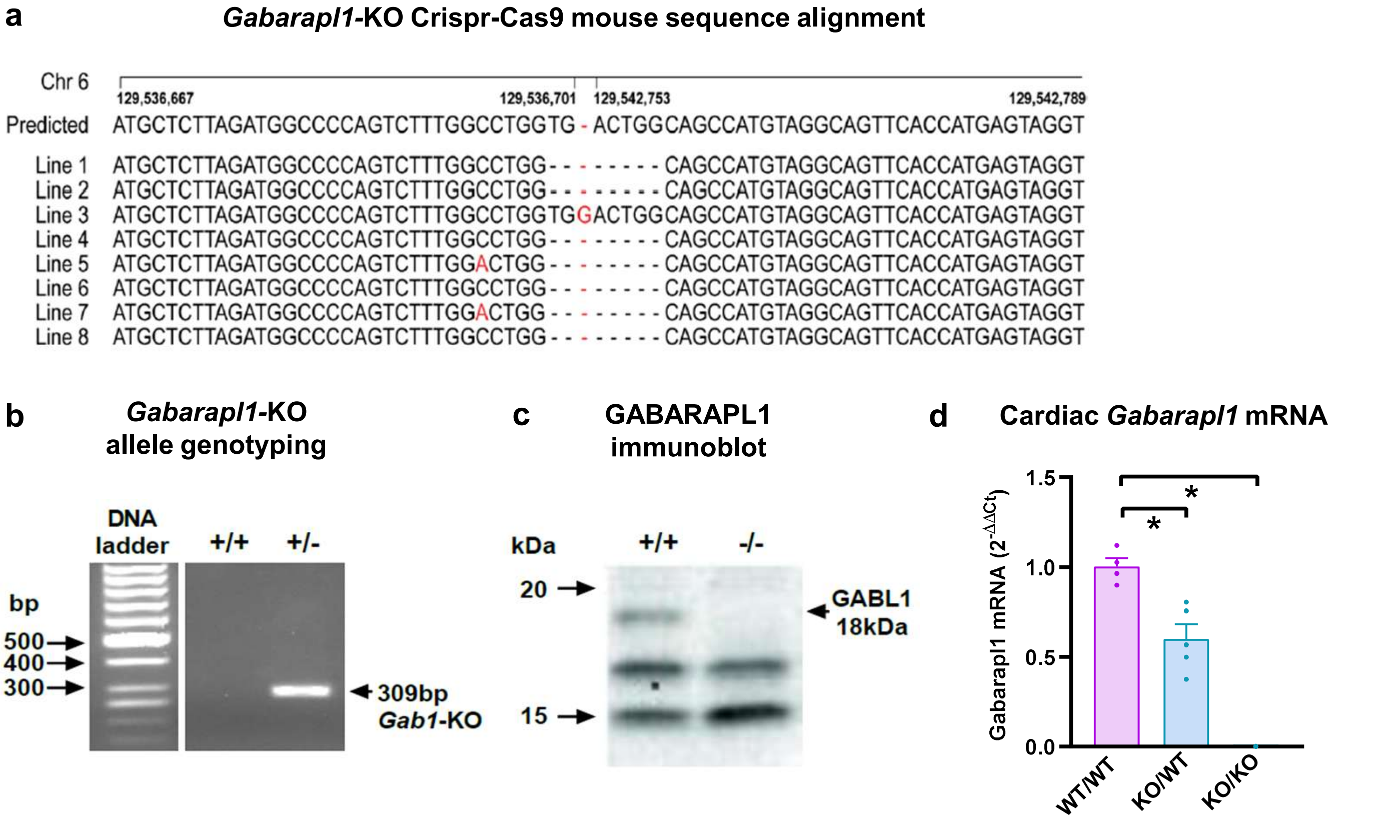


**Extended Data Fig.2 | Crispr-Cas9 *Gabarapl1*-KO mouse validation. a,** *Gabarapl1* sequence details for the 8 founder Crispr-Cas9 mouse lines confirming excision of exons 2-4 from the *Gabarapl1* gene (Next Generation sequencing). Text in red highlights mutations differing from the predicted sequence. **b,** Validation of *Gabarapl1* knockdown using DNA electrophoresis to identify the presence of the Gabarapl1-KO allele in the heterozygote Gabarapl1-KO (tail sample). **c,** Immunoblot to confirm the absence of the GABARAPL1 band in the homozygote knockout mouse (membrane-enriched fraction of heart homogenate). **d,** Heterozygote *Gabarapl1*-KO mice exhibit ~50% knockdown of the *Gabarapl1* gene (qPCR) in the heart, and absence of the *Gabarapl1* gene is confirmed in the homozygote *Gabarapl1*-KO mouse heart (male mice, 30 weeks old, n=4-5 mice). Data presented as mean ± s.e.m. *p<0.05. Related to Figure 3.

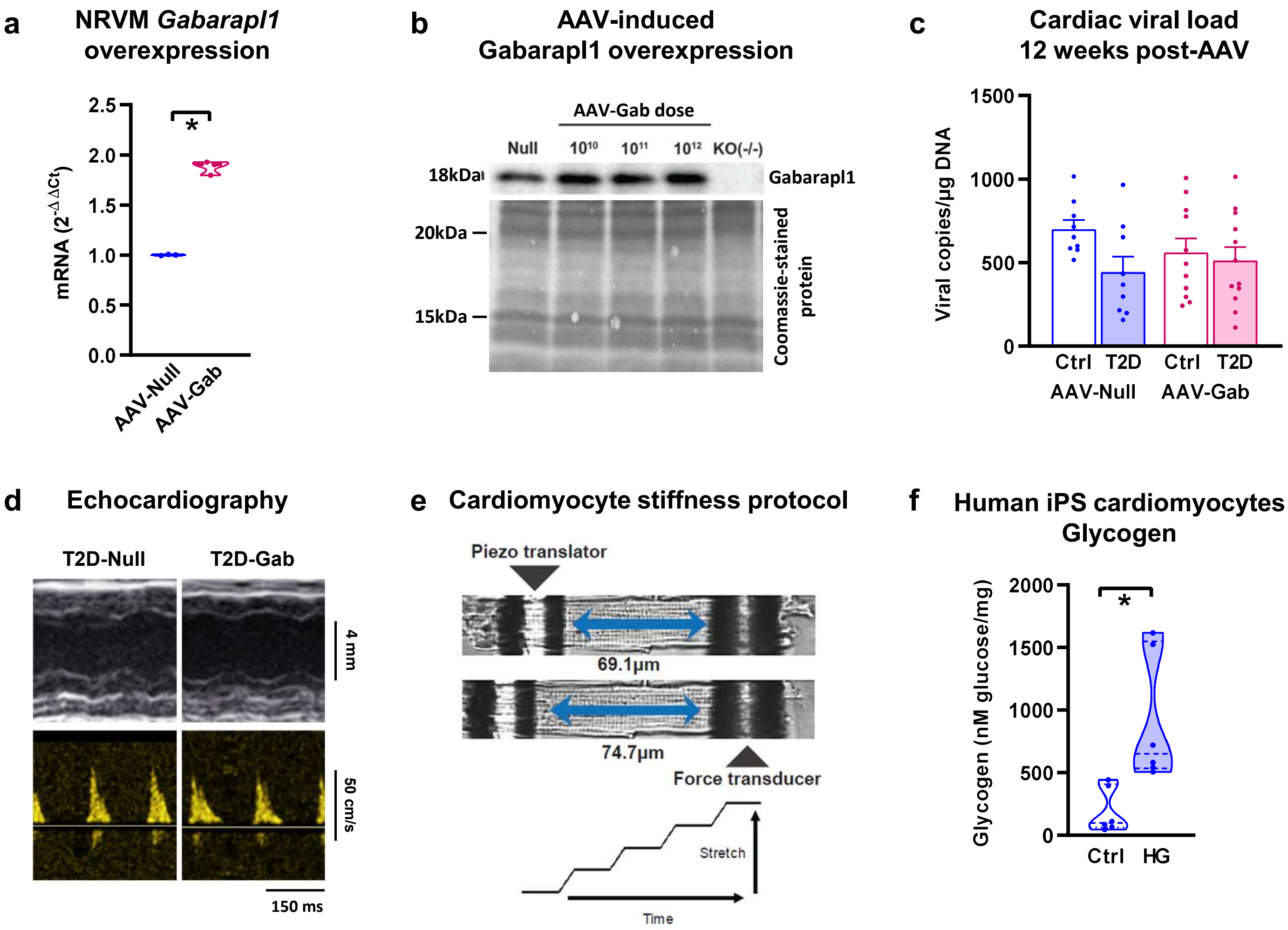


**Extended Data Fig.3 | AAV9-Gabarapl1 gene delivery *in vitro* and *in vivo*, cardiac stiffness protocol and iPS cardiomyocyte glycogen. a,** Confirmation of Gabarapl1 mRNA overexpression with AAV-Gabarapl1 in NRVMs (n=3 independent culture wells/group). **b,** Immunoblot of GABARAPL1 protein expression in mouse hearts 4 weeks post-injection (i.v.) of AAV-Null or AAV-Gab ($10^{10}$, $10^{11}$ or $10^{12}$ gc/mouse) with homozygote Gabarapl1-KO mouse heart as negative control. **c,** AAV vector burden in mouse heart 12 weeks post-injection (i.v.) of $10^{12}$ gc/mouse AAV9-cTnT-Null (AAV-Null) or AAV9-cTnT-Gabarapl1 (AAV-Gab) as measured by digital-droplet PCR detection of the WPRE viral element ($n \geq 9$ animals/group). **d,** M-mode and flow Doppler echocardiography exemplar traces in T2D mice treated with AAV-Null or AAV-Gab. **e,** Exemplar images of a non-stretched and stretched cardiomyocyte attached between two glass rods. Cardiomyocyte stretch protocol. **f,** Glycogen is increased in human iPS cardiomyocytes in response to high glucose exposure (n=6 wells/group). Data presented as mean ± s.e.m. *$p<0.05$. Related to Figure 4, 5 & 6.

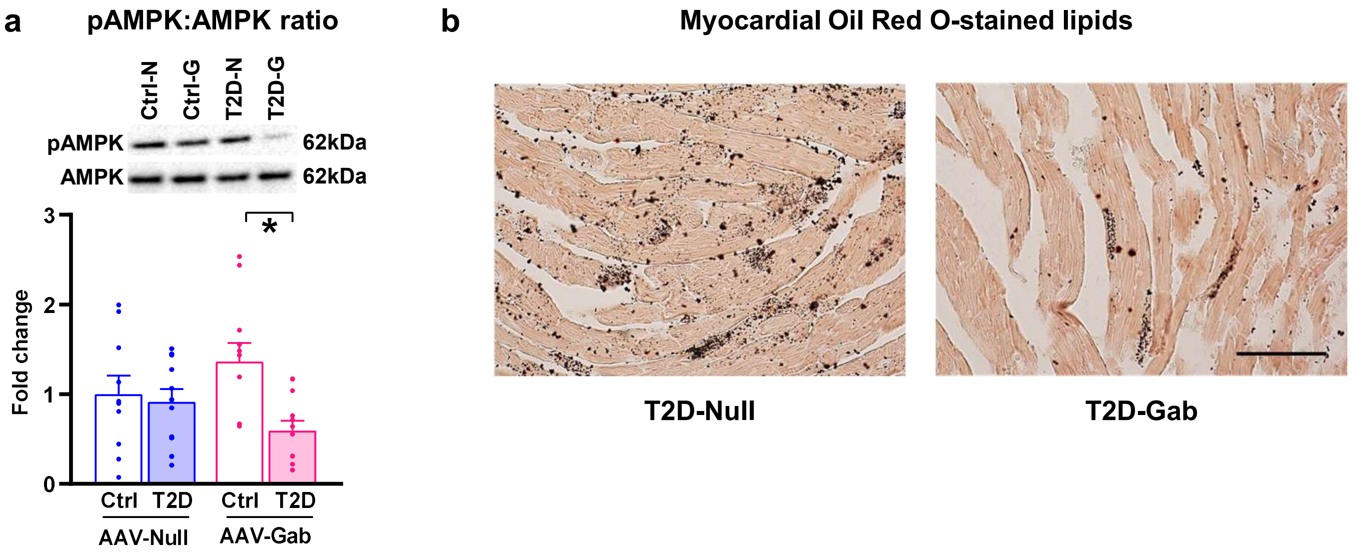


**Extended Data Fig.4 | AAV9-Gabarapl1 gene delivery is linked with AMPK signaling and myocardial lipid influence. a,** Ratio of phosphorylated (Thr172) to total AMPK protein expression is decreased with T2D in mice with Gabarapl1 gene delivery (AAV-Gab, n=6 animals/group). **b,** Myocardial sections from T2D mice with Gabarapl1 gene delivery (T2D-Gab) stained with Oil Red O for visualization of cellular neutral lipids (scale bar, 100μm).. Data presented as mean ± s.e.m. *p<0.05. Related to Figure 4.

**Extended Data Table 1 | Proteomic (LC-MS/MS) analysis of glycogen-associated proteins in cardiac and skeletal muscle in control and acute metabolic stress rats.** Differentially abundant and uniquely detected proteins from the Carbohydrate Metabolic Processes GO category in the glycogen proteome of cardiac and skeletal muscle from acute metabolic stress rats. In vivo 48hr hyperglycemia was induced by 55mg/kg i.p. streptozotocin destruction of pancreatic b-cells (n=4 rats/group). Related to Figure 1D.

| ACCESSION# | PROTEIN NAME | GENE NAME |
|---|---|---|
| CLH1_RAT | Clathrin heavy chain 1 | Cltc |
| CISY_RAT | Citrate synthase, mitochondrial | Cs |
| ENOG_RAT | Gamma-enolase | Eno2 |
| F16P2_RAT | Fructose-1,6-bisphosphatase isozyme 2 | Fbp2 |
| STBD1_RAT | Starch-binding domain-containing protein 1 | Stbd1 |
| LGUL_RAT | Lactoylglutathione lyase | Glo1 |
| ALAT1_RAT | Alanine aminotransferase 1 | Gpt |
| GLYG_RAT | Glycogenin 1 | Gyg1 |
| GSTO1_RAT | Glutathione S-transferase omega-1 | Gsto1 |
| HXK1_RAT | Hexokinase-1 | Hk1 |
| IDHC_RAT | Isocitrate dehydrogenase [NADP] cytoplasmic | Idh1 |
| MDHC_RAT | Malate dehydrogenase, cytoplasmic | Mdh1 |
| MTOR_RAT | Serine/threonine-protein kinase mTOR | Mtor |
| PFKAL_RAT | ATP-dependent 6-phosphofructokinase, liver type | Pfkl |
| PGAM1_RAT | Phosphoglycerate mutase 1 | Pgam1 |
| PGK1_RAT | Phosphoglycerate kinase 1 | Pgk1 |
| KPB1_RAT | Phosphorylase b kinase regulatory subunit alpha, skeletal muscle isoform | Phka1 |
| PHKG1_RAT | Phosphorylase b kinase gamma catalytic chain, skeletal muscle/heart isoform | Phkg1 |
| PK3CA_RAT | Phosphatidylinositol 4,5-bisphosphate 3-kinase catalytic subunit alpha isoform | Pik3ca |
| PP1B_RAT | Serine/threonine-protein phosphatase PP1-beta catalytic subunit | Ppp1cb |
| DHSO_RAT | Sorbitol dehydrogenase | Sord |
| TPIS_RAT | Triosephosphate isomerase | Tpi1 |

**Extended Data Table 2 | Characteristics of T2D mouse model.** Related to Figure 2.

| | Body weight (g) | Blood glucose (mM) | Glucose tolerance (area under GTT curve) |
|---|---|---|---|
| Control | 31.6 ± 1.3 | 10.7 ± 1.0 | 1924 ± 74 |
| T2D | 44.4 ± 1.4* | 14.6 ± 1.1* | 3363 ± 86 |

T2D, type 2 diabetes induced by high fat diet. Data analysed by Student's T-Test, *p<0.05. Data presented as mean ± SEM.

**Extended Data Table 3 | Primer sequences for glycophagy markers**. Related to Figure 2.

| Gene | Primer sequence 5' to 3' |
|---|---|
| Stbd1 | Forward: AAGCAGAGCATCTTCGAGAAAGC |
| | Reverse: ACCCAGTCTGCTCCAACATTC |
| Gabarapl1 | Forward: GGTCATCGTGGAGAAGGCTC |
| | Reverse: TAGAACTGGCCAACAGTGAGG |
| Gaa | Forward: CCTCGCAAGGTACCAACCTCTAC |
| | Reverse: GCAGGATGACATCCATGGCATTGC |
| 18S (housekeeper gene) | Forward: TCGAGGCCCTGTAATTGGAA |
| | Reverse: CCCTCCAATGGATCCTCGTT |

**Extended Data Table 4 | Patient characteristics.** Related to EDF1.

| | Non-diabetic | Type 2 diabetic |
|---|---|---|
| Sample size | 12 | 12 |
| Age (years) | 68 ± 6 | 66 ± 9 |
| Sex | 1F + 11M | 2F + 10M |
| Beta blockers | 8/12 | 10/12 |
| ACE inhibitors/Angiotensin Receptor Blockers | 7/12 | 9/12 |
| Statins | 6/12 | 10/12 |
| Metformin | 0/12 | 5/12 |
| Insulin | 0/12 | 7/12 |
| Hypertension | 7/12 | 9/12 |

**Extended Data Table 5 | siRNA sequences for *Gabarapl1* knockdown *in vitro*.** Related to Figure 2.

| | siRNA sequence (5'-3') | Genic location |
|---|---|---|
| Seq1 | UUCCCGUAGACACUUUCAU | Exon4: protein coding |
| Seq2 | AUUGCGAACAGCCCUAUUU | Exon4: UTR |
| Seq3 | UUUAACGCCAUCCAAACUG | Exon4: UTR |
| Seq4 | UCGCUUUGCAUUCCAGUGC | Exon4: UTR |

**Extended Data Table 6 | Guide RNA sequences for *Gabarapl1*-KO CRISPR-Cas9 mouse**. Related to Figure 3.

| | Spacer sequence (5'-3') | Strand | PAM (5'-3') | Genic location | Genomic location |
|---|---|---|---|---|---|
| gRNA1 | ggtggtgcgtcaaactatcg | + | cctggtg | Intron | Chr6:129,536,697 |
| gRNA2 | ggtctggtcccagatttgac | + | actggc | Post-gene | Chr6:129,542,735 |

**Extended Data Table 7 | Absolute glycogen values**. Related to Figure 4.

| | Glycogen (nmol glycosyl units/ mg protein) | | | |
|---|---|---|---|---|
| | Ctrl-Null | HG-Null | Ctrl-Gab | HG-Gab |
| NRVMs | 205 ± 10 | 311 ± 12* | 224 ± 17 | 243 ± 10 |

**Extended Data Table 8 | AAV9-Gabarapl1 T2D mice: systemic and cardiac characteristics.** Related to Figure 4&5.

| | AAV-Null | | AAV-Gab | |
|---|---|---|---|---|
| | Ctrl | T2D | Ctrl | T2D |
| Body weight (g) | 30.4 ± 0.7 | 46.1 ± 1.0* | 31.2 ± 0.7 | 46.5 ±1.4* |
| Blood glucose (mM) | 9.8 ± 0.8 | 12.3 ± 1.1* | 10.0 ± 0.5 | 11.9 ± 0.6* |
| Heart weight (mg) | 145.4 ± 6.0 | 169.1 ± 8.0* | 142.2 ± 3.0 | 171.4 ± 5.6* |
| Heart weight/body weight (mg/g) | 4.78 ± 0.1 | 3.68 ± 0.2* | 4.58 ± 0.1 | 3.70 ± 0.1* |
| Tibia length (mm) | 18.3 ± 0.1 | 18.1 ± 0.1 | 18.2 ± 0.1 | 18.3 ± 0.1 |
| Heart weight/tibia length (mg/mm) | 7.95 ± 0.3 | 9.32 ± 0.4* | 7.82 ± 0.1 | 9.35 ± 0.3* |
| Ejection fraction (%) | 70.7 ± 2.0 | 69.4 ± 1.6 | 66.3 ± 0.9 | 62.8 ± 2.2 |
| Fractional shortening (%) | 31.8 ± 1.1 | 29.8 ± 1.0 | 29.6 ± 0.6 | 26.8 ± 1.0 |

T2D mice (high fat diet-fed) injected with AAV9-cTnT-Null (AAV-Null) and AAV9-cTnT-Gabarapl1 (AAV-Gab) were assessed at 12 weeks post-AAV injection (i.v.) with 28 week diet feeding (n=10 mice/group). Data presented as mean ± SEM. *$p<0.05$.
